# Pain-regulation circuitry as a predictor of chronic pain phenotypes

**DOI:** 10.64898/2026.01.20.699535

**Authors:** Alireza Aleali, Muhammad Ali Hashmi, Alon Friedman, Ian Beaupre, Douglas Cane, Javeria Ali Hashmi

**Affiliations:** Department of Medical Neuroscience, Dalhousie University, Halifax, Nova Scotia, Canada; Department of Anesthesia, Pain Management, and Perioperative Medicine, Dalhousie University, Nova Scotia Health Authority, Halifax, NS, Canada; Massachusetts Institute of Technology, Cambridge, MA, United States; Departments of Cognitive and Brain Sciences, Physiology and Cell Biology, Ben-Gurion University of the Negev, Beer Sheva, Israel

**Keywords:** Chronic pain phenotyping, Fibromyalgia, Expectation-Induced Pain Modulation, Neuroimaging, Periaqueductal gray, Machine learning classification

## Abstract

Chronic pain involves sensory, emotional, and functional disruption, yet diagnostic labels rarely capture this variability, limiting individualized care. Multidimensional models, such as fear-avoidance and predictive coding, suggest that high emotional burden may disrupt expectation-based pain modulation and midbrain pain regulatory pathways, particularly within the periaqueductal gray (PAG). We tested whether chronic pain phenotypes defined by pain intensity, disability, and affective distress (PDA) differ in expectation-induced pain modulation, PAG connectivity, and behavioral markers including catastrophizing, hypervigilance, and medication use, and whether these features aid phenotype classification. We studied 159 patients with fibromyalgia or chronic low back pain and 72 controls. Our data-driven clustering approach identified high and low PDA groups. High PDA patients showed impaired modulation when positive expectations were violated and reported greater cognitive and behavioral burden (P<0.05). They also exhibited more negative connectivity between the dorsolateral/lateral PAG (dl/lPAG) and the dorsomedial prefrontal cortex (dmPFC) (corrected). Machine learning models classified PDA subtypes above chance, with accuracy improving when PAG connectivity was included. Findings highlight disrupted expectation-driven regulation and altered PAG pathways as markers for chronic pain stratification.

## Introduction

Chronic pain is a complex condition that includes sensory, affective, and cognitive dimensions, with considerable variability in symptoms and treatment response across patients, complicating diagnosis and treatment.^1–3^ Despite advances in understanding both peripheral and central mechanisms, substantial heterogeneity in clinical presentation and treatment response highlights the need for improved phenotyping to inform targeted interventions.^4,5^ The biopsychosocial framework supports the notion that individuals with high levels of Pain intensity, Disability, and Affective distress (PDA) may represent a distinct clinical phenotype with specific mechanistic underpinnings and therapeutic needs.^6–13^ Notably, this burden varies widely among individuals with chronic pain, and conventional diagnostic categories based on tissue pathology or syndromic labels, such as fibromyalgia (FM) or chronic low back pain (CLBP), may fail to capture this variability or inform treatment decisions.^3^ Classical evidence on pain modulation pathways centered on the periaqueductal gray area^14,15^ combined with new studies on pain expectation mechanisms^16–23^ suggests that abnormalities in these systems may represent a linchpin mechanism in exacerbating chronic pain symptoms^24,25^ and may contribute to recalcitrant refractory pain.^26^ However, these mechanisms remain insufficiently characterized and require further investigation to enable precision-based clinical stratification. This study aimed to identify a chronic pain phenotype characterized by high pain intensity, disability, and affective distress, to examine its behavioral and neurobiological features, and to evaluate its potential for improving diagnosis using machine learning.

Understanding how neurological pathways contribute to chronic pain is important because the brain integrates past and current experiences to predict incoming sensory information and shapes pain experiences. Predictive processing adjusts sensory input to align actual experiences with expectations^27^, and updates expectations based on sensory feedback to support safe adaptive interaction with the environment.^28–30^ In chronic pain, increased fear and hypervigilance can bias predictions to disrupt the normal integration of sensory input and expectations, resulting in amplified pain responses.^31–33^ Neurobiological alterations contribute to maladaptive avoidance behaviors, which, when reinforced over time, contribute to pain persistence, functional decline, and increased disability.^34–36^ Proper identification and intervention targeting these patterns are critical for proper therapy.^37,38^ However, no studies to date have directly compared Expectation-Induced Pain Modulation (EIPM) or its underlying mechanisms between individuals with chronic pain and healthy controls, nor examined its role in differentiating chronic pain phenotypes.

The periaqueductal gray (PAG) is a key hub in the descending pain modulatory system, integrating threat-related signals and regulating nociceptive input.^39–41^ Functionally distinct PAG subregions, the dorsolateral/lateral PAG (dl/lPAG) and ventrolateral PAG (vlPAG), are associated with active (sympathetic) and passive (parasympathetic) coping responses to physical threat, respectively.^39,42,43^ Disruptions in brain pathways connecting to these columns and especially the dl/lPAG pathways may impair descending inhibition and prediction error processing, contributing to chronic, treatment-resistant pain.^20,25,44,45^ Despite these roles, no studies to date have examined PAG subregion connectivity for phenotyping individuals with chronic pain.

The main objectives of this study were to (1) identify chronic pain phenotypes using cluster analysis based on pain intensity, disability, and affective burden (PDA axes); (2) determine whether PDA represents a specific clinical target by examining the role of fear avoidance behaviors and treatment refractoriness; (3) evaluate whether high PDA levels predict abnormal responses to expectation-related prediction errors and whether these abnormalities are associated with altered columnar PAG resting-state functional connectivity (rsFC); and (4) develop phenotype classifiers using a tiered approach, beginning with readily available features and assessing whether the addition of resource-intensive neuroimaging data improved classification accuracy.

## Materials and methods

### Study participants

Participants included healthy controls (*n* = 72), individuals with CLBP only (*n* = 109), CLBP and FM (*n* = 23) and those with FM only (*n* = 27). Recruitment was carried out through the Pain Management Unit (PMU) at the Queen Elizabeth II Health Sciences Centre, as well as through community advertisements, social media postings, and notices placed around Dalhousie University and the Victoria General Hospital in Halifax. All participants were required to be right-handed, between the ages of 18 and 75 years, and fluent in English to ensure comprehension of study procedures. All provided written informed consent prior to participation.

CLBP participants were included if they had experienced low back pain for six months or longer and reported an average of at least 4/10 clinical pain on the Brief Pain Inventory (BPI) during the two weeks preceding enrollment.^46,47^ FM participants were included if they met the American College of Rheumatology (ACR)^48^ diagnostic criteria for fibromyalgia, which require that (1) the Widespread Pain Index (WPI) be ≥7 with a Symptom Severity (SS) score ≥5, or that WPI range from 3–6 with an SS score ≥9; (2) symptoms be present at a similar level for at least three months; and (3) no other disorder better explain the pain. All FM participants had a confirmed clinical diagnosis established by a pain specialist, whereas 32 of the CLBP participants were included based on self-reported clinical history. Healthy controls were included if they reported no ongoing acute or chronic pain, no neurological or systemic medical disorders, and no use of analgesic or psychotropic medications. Exclusion criteria applicable to all groups included a history of significant cardiac, respiratory, or neurological disorders; contraindications to MRI scanning (e.g. pacemaker, metallic implants, claustrophobia, or pregnancy); uncorrected visual impairments; or active participation in a treatment program for pain relief within the previous two months.

### Identifying chronic pain phenotypes through symptom-based stratification

To identify PDA phenotypes, standardized scores from three categories of measures were analyzed using Principal Component Analysis (PCA)^49,50^ to extract latent components that captured core dimensions of symptom burden. The scores included pain severity and disability measured using the BPI questionnaire. Depression (Beck Depression Inventory (BDI))^51^ and state and trait anxiety (the Spielberger Trait Anxiety Inventory (STAI)) ^52^ for affective distress. The retained principal components were then entered into a two-stage clustering pipeline. First, hierarchical clustering^53^ was used to determine the optimal number of clusters, followed by K-means clustering^54^ for final group assignment. The resulting groups were subsequently age- and sex-matched to healthy controls (see Supplementary section 2).

The identified phenotypes were characterized for differences in clinical symptoms with the McGill Pain Questionnaire (MPQ)^55^, the Pain Vigilance and Awareness Questionnaire (PVAQ)^56^, and the Pain Catastrophizing Scale (PCS).^57^ Medication use was quantified on the Medication Quantification Scale (MQS).^58^ The number of over-the-counter (OTC) medications and prescribed medications were assessed to check for pharmacological treatment refractoriness. All questionnaires were administered via REDCap (http://www.project-redcap.org).

### Test for expectation-induced pain modulation (Schema task)

Participants were evaluated on an expectation task (schema task), which estimates the interplay between top-down cognitive expectations and bottom-up sensory inputs in modulating pain perception. By manipulating the relationship between visual cues and thermal stimuli, task conditions are created where participants’ expectations are either matched or mismatched to the noxious thermal stimulus, and the resulting pain ratings are assessed to observe how the mismatch (prediction errors) interact to shape pain experiences.^59^

In total, 216 participants completed the task, including 65 HC, 102 individuals with CLBP, and 49 individuals with FM of which 60% of tests were conducted inside the scanner. Participants were informed that they would be shown visual information on a screen, experience thermal stimuli (heat), and subsequently rate the pain intensity they perceived using a numeric scale. During each trial, participants were presented with the threat cues on the screen, which either predicted the upcoming stimulus intensity as a threat range (e.g. “the incoming heat stimulus is at 70%–80% intensity”) in the matched run or as a threat value (e.g. “the incoming heat stimulus is at 80% intensity”). Following the cue, a red fixation cross was displayed on the screen and heat stimuli were delivered simultaneously to their skin on the lower left leg (specifically, the tibialis anterior muscle). Stimulation was delivered using the PATHWAY system (Medoc Ltd., Ramat Yishai, Israel) in 63% of healthy controls and 26% of FM patients; all remaining participants were tested using the TCS-II system (QST.Lab, Strasbourg, France). Participants then rated their perceived pain intensity on a 0–100 numerical rating scale (NRS), where 0 indicated "no pain at all" and 100 represented "the worst pain imaginable." They used a response pad (Lumina LSC-400 controller, Cedrus) or a keyboard to move a cursor along the scale and select the number corresponding to their pain level (Figure 1A). This procedure was repeated multiple times across four different runs, including one matched-condition run and three mismatched-condition runs. To assess potential effects of stimulation equipment on pain ratings, a multivariate analysis of variance was performed on ranked ratings across cue levels, with equipment type as the main factor and age and sex as covariates. Equipment type had no significant effect on overall pain ratings (*P* = 0.25).

**Figure 1.**
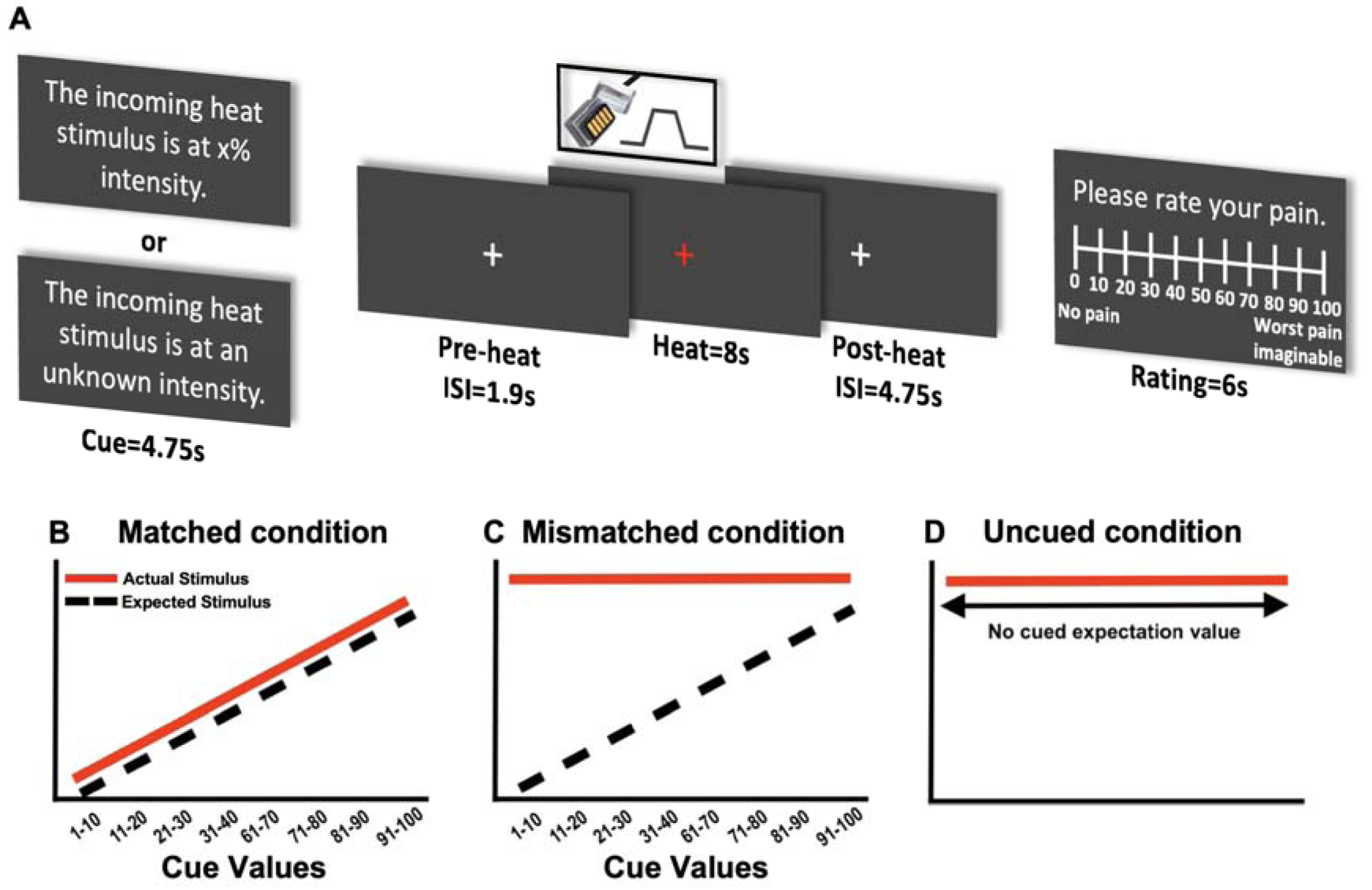
Experimental Paradigm of the Schema Task for Pain Perception. ***A,*** In each trial, participants first viewed a threat cue on the screen indicating the intensity of the upcoming heat stimulus. Subsequently, the heat pain stimulus was administered while a red fixation cross was presented. Following the heat stimulus, participants were asked to rate their pain intensity on a scale from 0 to 100. ***B*,** Matched condition: The threat cue predictions corresponded directly to the heat stimulus intensity linearly in this condition. This design allowed participants to develop a linear schema for perceived pain, where the actual heat stimulus intensity was consistent with their expectations based on the visual cues. ***C,*** In the first mismatched run (Mismatched Level 1), cue values from 1–40% were paired with low-heat (45°C) stimuli, while cue values from 61–100% were paired with high-heat (47°C) stimuli, creating prediction errors of 0°C to 1.2°C. In the following runs (Mismatched Level 2), cue values from 1–40% were again paired with 45°C stimuli, while cue values from 1–100% were paired with 47°C stimuli, resulting in prediction errors of 0°C to 3.2°C. In both levels, there was no prediction error for cues in the 91–100% range. ***D,*** Uncued condition: no cued expectation value was provided, and the temperature was kept constant at 45°C and 47°C (red line). This condition was used to assess pain perception without any prior expectation.

In the matched run (Figure 1B), each threat cue range and stimulus pairing was designed so that the cues accurately predicted the pain intensities and the association was linear, where each 10-point increase in the cued threat value corresponded to a 0.4°C increase in stimulus temperature within a 43.8°C to 47°C range. In this way, participants learned to anticipate the pain intensity based on the threat cues.

In the Mismatched Level 1 condition (second run, following the matched run), cue values from the range of 1–40% were paired with low-heat stimuli (45°C), while cue values from 61–100% were paired with high-heat stimuli (47°C). This design created a linear increase in the difference between the expected and actual temperatures for the cue ranges of 1–40% and 61–100%, with the prediction error varying between 0°C and 1.2°C. There was no prediction error for cue ranges 31–40% and 91–100%, where the expected and actual heat intensities matched. In the Mismatched Level 2 condition (third and fourth runs), cue values from 1–40% were again paired with 45°C stimuli, maintaining a low range of prediction errors. However, in these runs, cue values ranging from 1 to 100% were paired with a constant 47°C stimulus, introducing a wider range of prediction errors (0–3.2°C). Similar to the first mismatched run, no prediction error occurred for cues in the 91–100% range, where the expectation matched the actual stimulus (Figure 1C).

Additionally, in the uncued condition (Figure 1D), no threat cues were provided. For the control condition (bottom-up only), the cues showed that “the incoming heat stimulus intensity is unknown”. Heat stimuli in this condition were delivered at two fixed intensities (47°C and 45°C). Across all four runs, each cue level (e.g. 21-30) was repeated a minimum of 4 times and a maximum of 7 times. Additionally, there were 6 trials of the uncued condition at 47°C and 10 trials at 45°C. Additionally, to quantify the effects of both top-down (TD) and bottom-up (BU) processes on pain modulation, specific metrics were calculated for each participant.

#### Bottom-Up prediction error responses

Bottom-up prediction error represents the response to the violation of positive expectations (low threat cues) and was measured by taking the difference in pain ratings between matched and mismatched temperatures that were paired with the same range of cues (1-90). Since the cues were identical in both conditions, the prediction errors were driven by a combination of bottom-up sensory sources and prediction error. A higher difference in pain ratings in response to mismatched temperatures indicated a higher pain sensitivity to bottom-up prediction errors.

Specifically, for each participant, the metric was computed as:

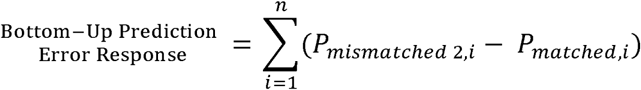

Where:

*P_mismatched_*_2,*i*_ represents the pain rating in the mismatched condition level 2 for cue level *i*, and *P_matched_*_,*i*_ represents the pain rating in the matched condition for the same cue level *i*. *n* represents the number of cue ranges considered (in this case, 7 cue ranges: 1–10, 11–20, 21–30, 31–40, 61–70, 71–80, and 81–90). The cue range 91–100, the visual threat cues in this range still correspond to the highest heat stimulus (47°C) had no predictor error, hence excluded from calculation.

To assess the maximum effect of violations in the positive cues, we computed the prediction error max response. This metric was calculated as the difference in pain ratings, which occurs when the lowest visual threat cues (1–20) are paired with heat stimuli of 45°C and 47°C in the mismatched conditions. This metric was calculated as:

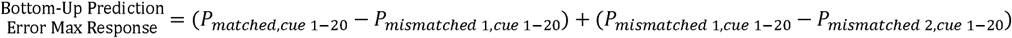

Where:

*P_matched, cue_*_1-20_, *P_mismatched_* _l*, cue*1-20_, and *P_mismatched_* _2*, cue*1-20_ are the average pain ratings for the cue range 1–20% in the matched, mismatched 1 (45°C), and mismatched 2 (47°C) conditions, respectively.

#### TD prediction error response

As previously described^60^, the TD effect was assessed for events where the mismatch resulted from violations in cues, but bottom-up temperatures were constant, so that the cues were maximally different, but the temperature was held at an identical temperature. This was determined by analyzing the difference in pain ratings between the maximum prediction error range: 1–20% and the minimum prediction error range: 80–100% while temperature was at the same intensity (47°C). Thus:

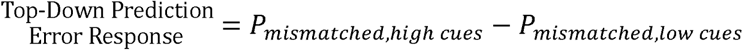

Where:

*P_mismatched_*_, *high cues*_ refers to the average pain rating for the highest threat cues (80–100%) in the mismatched condition, and *P_mismatched, low cues_* refers to the average pain rating for the lowest threat cues (1–20%) in the mismatched condition. In both cases, the temperature was kept at 47°C, but the visual threat cues were different.

### MRI data acquisition and preprocessing

Structural (T1) and an 8-minute resting state functional MRI (rsfMRI) scan were collected on a 3T scanner. Data preprocessing followed established pipelines using Analysis of Functional NeuroImages (AFNI; http://afni.nimh.nih.gov/afni) and FMRIB Software Library (FSL; http://www.fmrib.ox.ac.uk/fsl), incorporating motion correction, distortion correction, spatial smoothing, nuisance regression, and registration to standard space. Participants with excessive motion or DVARS (difference in signal intensity between successive volumes) outliers were excluded, and motion metrics were included as nuisance covariates in subsequent analyses to control for residual artifacts. Main findings were tested for effects of smoothing, and by using a whole brain stem ROI (region of interest) as a control region. Since heart rate and respiration data were available in the patient groups, RETROICOR (RETROspective Image CORrection) was used for cleaning brainstem artifact, and the effect of artifacts on main findings was established only in the patient groups. For full details of acquisition and preprocessing see supplementary section 3.

#### Time series extraction from parcellations

Time series were extracted from parcellations based on the Optimized Harvard-Oxford scheme. ^61,62^ For each participant, blood oxygenation level-dependent (BOLD) time series were extracted from each voxel and averaged for each region. The same procedure was applied to the PAG subregions and the whole brainstem, as defined by the parcellation. (See Supplementary Table 1 for full parcellation details and supplementary section 4 for PAG characteristics).

#### Resting state functional connectivity (rsFC) analyses

##### Seed-Based connectivity analysis

This analysis builds upon our previous work^39^, where we examined whole brain rsFC differences between dl/lPAG and vlPAG in a smaller sample size of healthy controls. Prior to analysis, participants with missing resting-state fMRI data were excluded, and the remaining sample was age- and sex-matched across the three study groups to control for potential confounds (see Supplementary Figure 1 for a flowchart of participant inclusion and exclusion, including details on age- and sex-matching). We then investigated whether the dl/lPAG and vlPAG exhibit distinct connectivity profiles by comparing their rsFC within each of the three groups, followed by group comparisons. For each participant, the right and left dl/lPAG and vlPAG were considered separately, and 0-lag Pearson correlation coefficients^63^ were computed with 134 predefined brain regions, generating 4 × 134 adjacency matrices. Given the non-normal distribution of the data, the resulting connectivity matrices were analyzed for within-subject differences between dl/lPAG and vlPAG using Wilcoxon signed-rank tests. To examine between-group differences in absolute rsFC for each PAG subregion, Mann–Whitney U tests were used to perform all pairwise comparisons. False discovery rate (FDR) correction (*q* < 0.05) was applied across all tests to control for multiple comparisons.

### Data analyses

We first assessed continuous variables for missing values and tested for normality using the Kolmogorov–Smirnov and Shapiro–Wilk tests. When assumptions of normality were met, parametric tests (e.g. t-tests, one-way ANOVA) were used. When assumptions were violated, non-parametric alternatives were applied (e.g. Mann–Whitney U test, Kruskal–Wallis test). Within-group comparisons were analyzed using rank-transformed repeated-measures ANOVA to accommodate non-normal distributions while preserving the factorial structure. When more than two within-subject conditions were compared, non-parametric tests were used, with Bonferroni correction for post hoc pairwise comparisons. Categorical variables were analyzed using chi-square tests. Multiple comparisons were corrected using FDR adjustment for connectivity analyses. To reduce potential confounding, groups were age- and sex-matched prior to all analyses. In addition, age and sex were also tested as covariates in the rsFC analyses.

Classification accuracy was evaluated using Receiver Operating Characteristic (ROC) curves, logistic regression, and machine learning methods. Statistical analyses were conducted in IBM SPSS Statistics (Version 29.0.2), MATLAB R2024b (MathWorks Inc., Natick, MA), and RStudio (Version 2024.09), which was also used for data visualization.

### Modeling chronic pain phenotypes using stepwise supervised machine learning

To classify chronic pain severity using domain-informed features, a stepwise supervised machine learning approach was applied across three models: (1) clinical/behavioral data alone, (2) clinical plus schema-based pain modulation metrics, and (3) all prior features plus rs-fMRI connectivity. Standard preprocessing included missing value imputation, numeric casting, and Lasso regularization for dimensionality reduction. Highly correlated features were reviewed for redundancy. Models were trained using a stratified 70–30 train-test split with five-fold cross-validation, and evaluated using logistic regression, random forest, and support vector machine (SVM). Performance was assessed on the test set via accuracy and weighted F1-score; Shapley Additive exPlanations (SHAP)^64^ analyses interpreted key predictors in the best model (for more details, see supplementary section 5).

## Results

### Derivation of pain–disability–affect phenotypes via principal component and cluster analysis

PCA was performed to reduce data dimensionality across pain impact and affective distress measures. Suitability tests confirmed its applicability (*KMO* = 0.691; *Bartlett’s test*: *χ*² = 354.90, *df* = 10, *P* < 0.001). Two components with eigenvalues >1 were extracted. After Varimax rotation, the first component retained high loadings on BDI (0.886), state anxiety (0.884), and trait anxiety (0.939), capturing affective distress. The second component showed strong loadings on BPI pain severity (0.912) and pain interference (0.873), indicating pain impact (Figure 2A,B). The distribution of FM, CLBP, and CLBP+FM participants showed substantial overlap across both PCA components, suggesting that diagnosis alone does not adequately capture symptom variability (Figure 2D).

**Figure 2.**
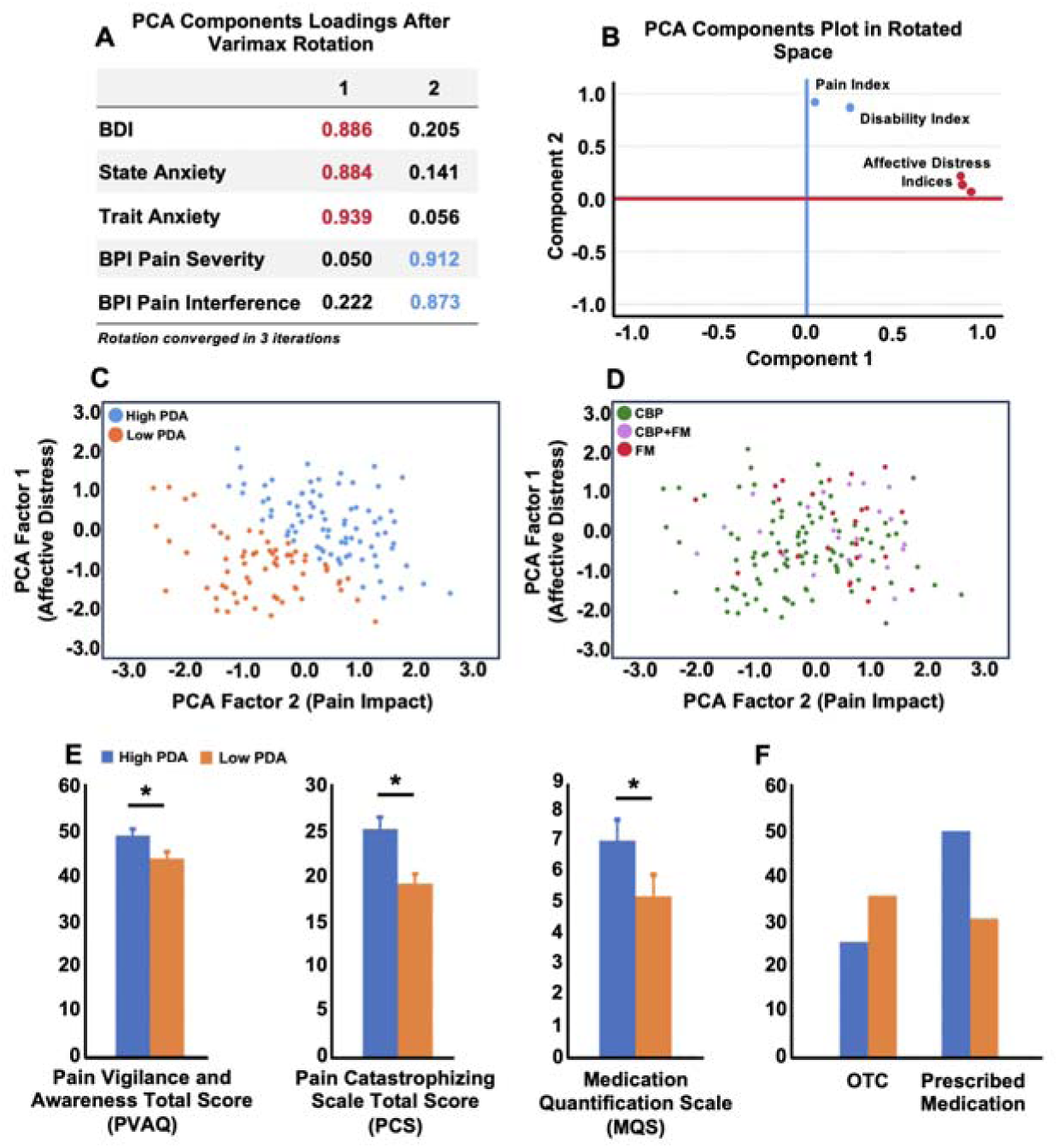
PDA phenotypes derived from principal component and clustering analyses. ***A,*** Rotated component matrix after Varimax rotation showing two factors: affective distress (loading on BDI, state and trait anxiety) and pain impact (loading on pain severity and pain interference). ***B,*** PCA component plot illustrating separation of affective distress and pain impact dimensions. ***C,*** K-means clustering in rotated PCA space differentiating high PDA (blue) and low PDA (orange) groups. ***D,*** Distribution of patients with chronic low back pain (green), fibromyalgia (red), or both (purple) across PCA space, showing diagnostic overlap. ***E,*** High PDA participants scored higher on the PVAQ, PCS, and MQS (p < 0.05). ***F,*** High PDA participants were more likely to use prescribed than OTC medications (Pearson’s χ², p = 0.021). BDI, Beck Depression Inventory; BPI, Brief Pain Inventory; CLBP, Chronic Low Back Pain; FM, Fibromyalgia; OTC, Over-the-Counter; PCA, Principal Component Analysis; PDA, Pain-Disability-Affect.

Individuals who loaded higher on the two principal components were grouped as high PDA phenotype based on hierarchical and k-means clustering (Figure 2C), and cluster validity was confirmed via silhouette analysis (for details, see supplementary section 2). Following cluster assignment based on chronic pain severity, participants were age- and sex-matched to form three comparable groups: high PDA, low PDA, and HC. The final behavioral sample consisted of 60 HC participants, 75 in the high PDA group (34 FM and 41 CLBP), and 65 in the low PDA group (13 FM and 52 CLBP).

High PDA individuals showed greater treatment refractoriness, reflected in high pain and distress despite a higher medication burden (MQS; *P* = 0.036) (Figure 2E), and a greater reliance on prescribed rather than over-the-counter medications. This pattern was supported by both a chi-square test of independence (*P* = 0.021) and a linear-by-linear association test (*P* = 0.018), indicating a graded shift toward prescription use across increasing severity levels. A summary of clinical and affective metrics is presented in Table 1.

**Table 1.**
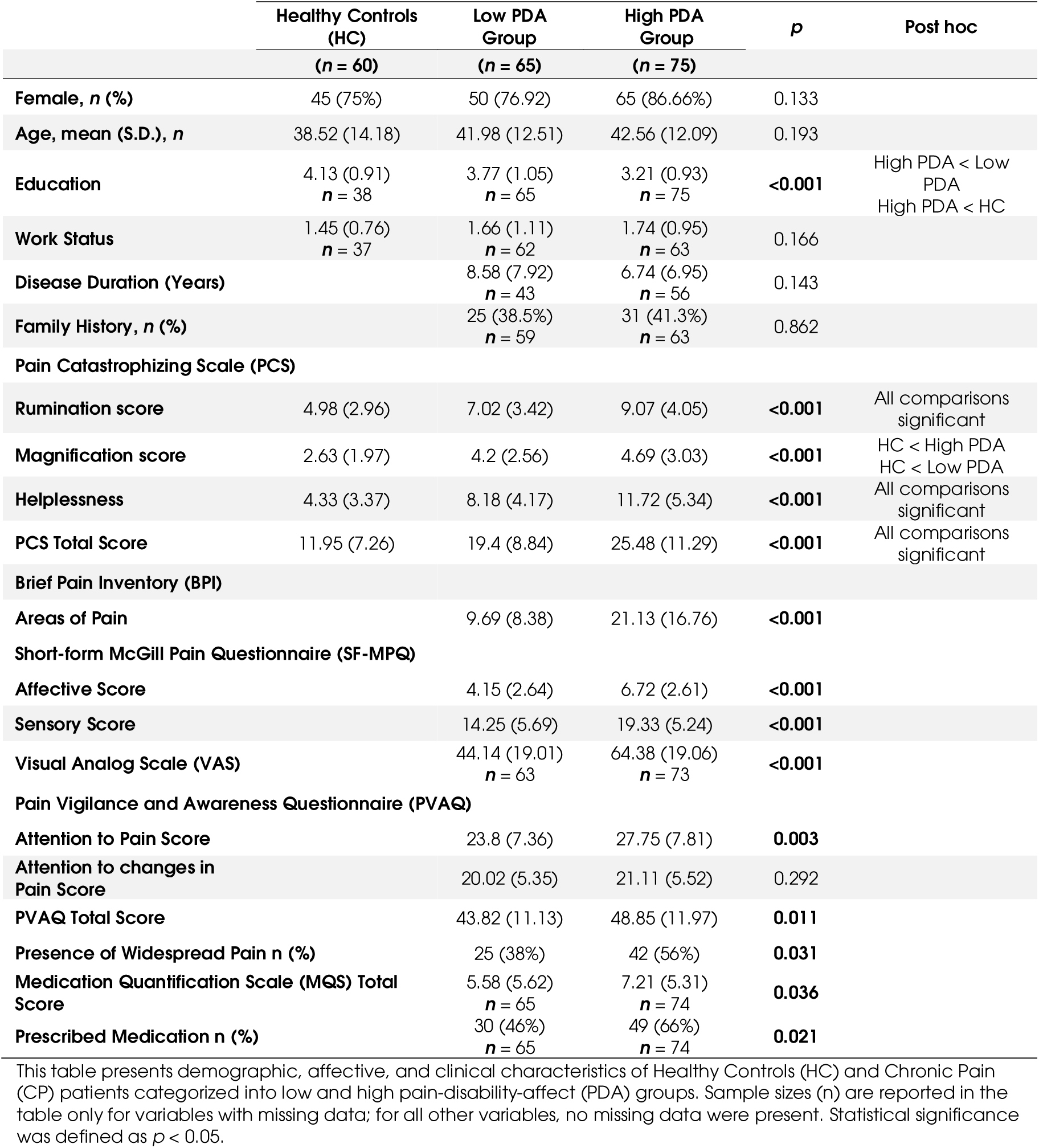
Behavioral and Clinical Distinctions of the High Pain–Disability–Affect Phenotype.

|  | Healthy Controls<br>(HC)<br>(*n* = 60) | Low PDA<br>Group<br>(*n* = 65) | High PDA<br>Group<br>(*n* = 75) | *p* | Post hoc |
| --- | --- | --- | --- | --- | --- |
| Female, *n* (%) | 45 (75%) | 50 (76.92) | 65 (86.66%) | 0.133 |  |
| Age, mean (S.D.), *n* | 38.52 (14.18) | 41.98 (12.51) | 42.56 (12.09) | 0.193 |  |
| Education | 4.13 (0.91)<br>*n* = 38 | 3.77 (1.05)<br>*n* = 65 | 3.21 (0.93)<br>*n* = 75 | <0.001 | High PDA < Low PDA<br>High PDA < HC |
| Work Status | 1.45 (0.76)<br>*n* = 37 | 1.66 (1.11)<br>*n* = 62 | 1.74 (0.95)<br>*n* = 63 | 0.166 |  |
| Disease Duration (Years) |  | 8.58 (7.92)<br>*n* = 43 | 6.74 (6.95)<br>*n* = 56 | 0.143 |  |
| Family History, *n* (%) |  | 25 (38.5%)<br>*n* = 59 | 31 (41.3%)<br>*n* = 63 | 0.862 |  |
| **Pain Catastrophizing Scale (PCS)** |  |  |  |  |  |
| Rumination score | 4.98 (2.96) | 7.02 (3.42) | 9.07 (4.05) | <0.001 | All comparisons significant |
| Magnification score | 2.63 (1.97) | 4.2 (2.56) | 4.69 (3.03) | <0.001 | HC < High PDA<br>HC < Low PDA |
| Helplessness | 4.33 (3.37) | 8.18 (4.17) | 11.72 (5.34) | <0.001 | All comparisons significant |
| PCS Total Score | 11.95 (7.26) | 19.4 (8.84) | 25.48 (11.29) | <0.001 | All comparisons significant |
| **Brief Pain Inventory (BPI)** |  |  |  |  |  |
| Areas of Pain |  | 9.69 (8.38) | 21.13 (16.76) | <0.001 |  |
| **Short-form McGill Pain Questionnaire (SF-MPQ)** |  |  |  |  |  |
| Affective Score |  | 4.15 (2.64) | 6.72 (2.61) | <0.001 |  |
| Sensory Score |  | 14.25 (5.69) | 19.33 (5.24) | <0.001 |  |
| Visual Analog Scale (VAS) |  | 44.14 (19.01)<br>*n* = 63 | 64.38 (19.06)<br>*n* = 73 | <0.001 |  |
| **Pain Vigilance and Awareness Questionnaire (PVAQ)** |  |  |  |  |  |
| Attention to Pain Score |  | 23.8 (7.36) | 27.75 (7.81) | 0.003 |  |
| Attention to changes in Pain Score |  | 20.02 (5.35) | 21.11 (5.52) | 0.292 |  |
| PVAQ Total Score |  | 43.82 (11.13) | 48.85 (11.97) | 0.011 |  |
| Presence of Widespread Pain *n* (%) |  | 25 (38%) | 42 (56%) | 0.031 |  |
| Medication Quantification Scale (MQS) Total Score |  | 5.58 (5.62)<br>*n* = 65 | 7.21 (5.31)<br>*n* = 74 | 0.036 |  |
| Prescribed Medication *n* (%) |  | 30 (46%)<br>*n* = 65 | 49 (66%)<br>*n* = 74 | 0.021 |  |

This table presents demographic, affective, and clinical characteristics of Healthy Controls (HC) and Chronic Pain (CP) patients categorized into low and high pain-disability-affect (PDA) groups. Sample sizes (*n*) are reported in the table only for variables with missing data; for all other variables, no missing data were present. Statistical significance was defined as $p < 0.05$ .

### Predictive cues modulate pain in high PDA, low PDA, and healthy controls

Previous research from our lab has demonstrated that predictive cues influence pain perception in healthy participants.^59^ In this study, we reproduce these findings by demonstrating this effect separately in all three groups, where cues with successively lower threat cues produced a linear reduction in pain even though the stimulus temperature was identical. The analysis examined matched and mismatched conditions within each group using a two-way repeated-measures ANOVA (matched vs. mismatched conditions × cued threats at eight levels). Results indicated significant effects in all groups: for the high PDA group, *F_(7,55)_* = 11.097, *P* < 0.001; for the low PDA group, *F_(7, 54)_* = 8.216, *P* < 0.001; and for the HC group, *F_(7, 46)_* = 13.456, *P* < 0.001). (Figure 3A–C).

**Figure 3.**
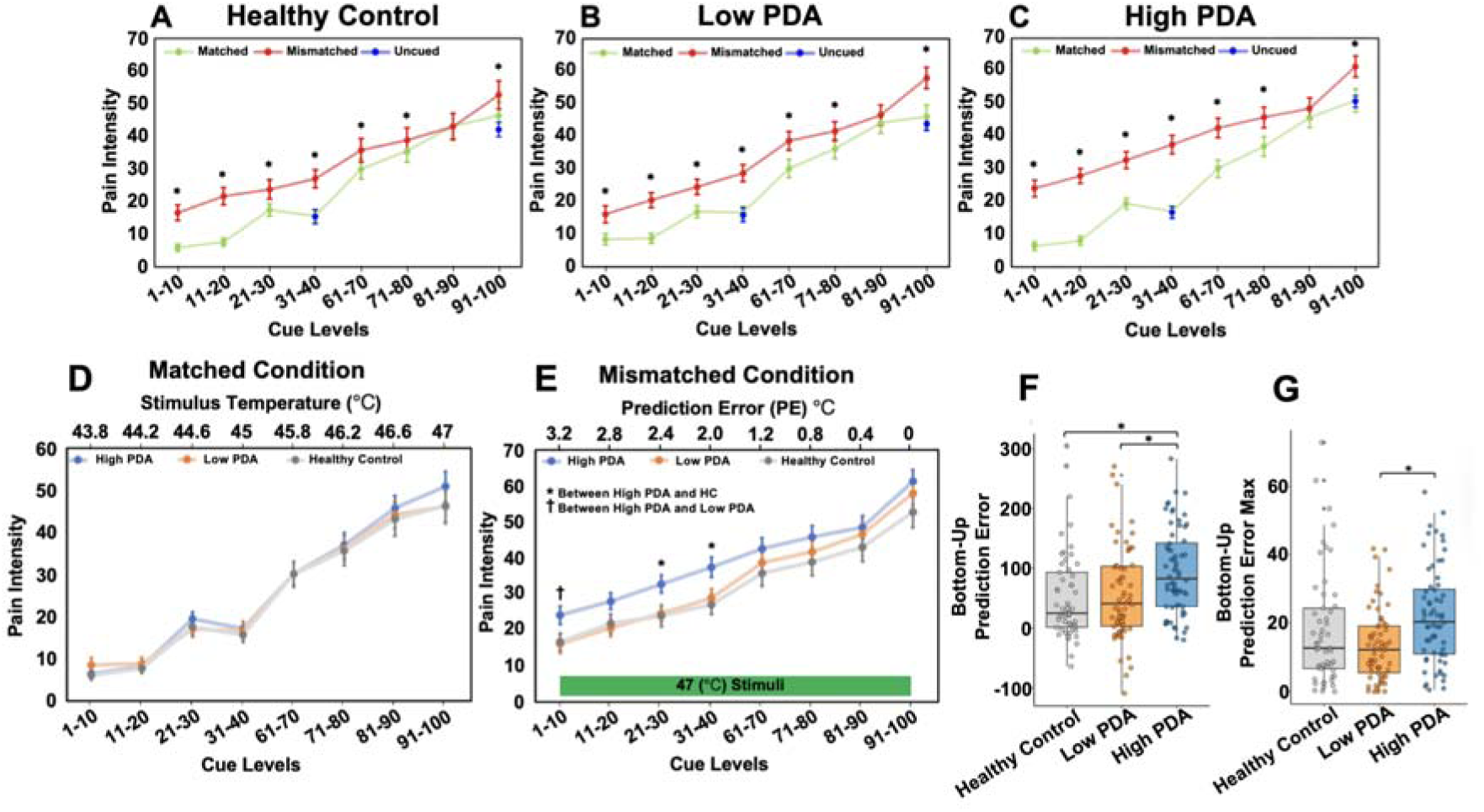
Expectation-driven modulation of pain perception across groups in the schema task. ***(A–C)*** Within-group comparisons of pain intensity ratings across cue levels in matched (green), mismatched (red), and uncued (blue) conditions for Healthy Controls (A), Low PDA (B), and High PDA (C). ***D,*** Matched Condition plot: The x-axis represents the cue levels, ranging from 0-10 to 91-100, and the y-axis represents the pain intensity ratings, showing pain intensity ratings change with change in cue-temperature pairings in all three groups. The blue line represents the high PDA, the orange line represents the low PDA, and the grey line represents the Healthy Controls. ***E,*** Mismatched Condition plot: showing varying pain intensity ratings despite a constant temperature (47°C), reflecting reliance on expectation cues. The high PDA group showed higher pain ratings at low threat cues compared to the other groups. Error bars indicate the ±SEM. Significant differences between groups are marked with asterisks (*p < 0.05) and dagger (†). ***F,*** Bottom-Up Predictor Error Response shown as box plots, which were determined by subtracting the pain ratings for each cue level in the mismatched condition from those in the matched condition and then summing these differences (excluding the 91-100 cue level, as it is matched in both conditions). ***G,*** Bottom-Up Predictor Error Response Max shown as a box plot, which is a composite metric derived from pain ratings at the lowest threat cue range (1–20%), calculated from differences between matched and mismatched (45°C and 47°C) conditions. PDA, Pain-Disability-Affect.

When pain ratings were compared across matched, mismatched, and uncued trials at 45°C and 47°C, both high and low PDA groups showed higher ratings in mismatched than uncued trials at both temperatures, with no difference between matched and uncued trials. In contrast, healthy controls showed higher ratings in mismatched than uncued trials only at 47°C, where matched trials also exceeded uncued ratings; no cueing effects were observed at 45°C (see Supplementary Section 6 for details).

### High PDA group demonstrates abnormalities in the expectation task

There were no significant group differences when threat cues accurately predicted pain intensity (matched condition; Figure 3D). In contrast, in the mismatched condition, the high PDA group reported significantly higher pain ratings than both the low PDA and HC groups, particularly for lower threat cues (1–40) (Figure 3E). These differences were significant for three of the four low-threat cues. The low PDA group, however, did not differ substantially from the HC group. Overall, the high PDA group showed the greatest pain ratings in response to violated positive expectations.

Thus, a three-way mixed ANOVA (three factors: (1) Condition (matched vs. mismatched), (2) Cued Threat Levels (eight levels: 1–10, 11–20, 21–30, 31–40, 61–70, 71–80, 91–100), and (3) Group (High PDA, Low PDA, and Healthy Control) as a between-subject factor) showed a significant interaction effect for Condition × Cued Threat Levels × Group interaction (*F_(14,336)_* = 2.227, *P* = 0.034). Additionally, there was a significant main effect for Cued Threat Levels (*F_(7,167)_* = 90.483, *P* < 0.001), and for Condition (matched vs. mismatched) (*F_(_*_1,173)_ = 158.032, *P* < 0.001), showing that both the cue levels and the condition (matched vs. mismatched) overall significantly influenced pain ratings.

Additionally, no significant group differences were observed in the uncued conditions at 47°C (*P* = 0.140) or 45°C (*P* = 0.541). See Supplementary Section 6 and Supplementary Figure 11 for further details.

Group differences were mainly driven by greater sensitivity to violated positive expectations gauged from bottom-up temperatures in the high PDA group. Both the bottom-up prediction error response and the bottom-up prediction error maximum response varied significantly across groups (*P* = 0.001 and *P* = 0.003, respectively; Figure 3F,G), with the high PDA group showing higher values than both the HC and low PDA groups. In contrast, the TD prediction error response did not differ significantly between groups (*P* = 0.416; Supplementary Figure 12).

Significant positive correlations were found between bottom-up prediction error response and PCS (*ρ* = 0.148, *P* = 0.035) and PVAQ (*ρ* = 0.198, *P* = 0.019), suggesting that greater pain hypervigilance predicts higher sensitivity to bottom-up prediction errors (see Supplementary Figure 13).

### PDA related abnormalities in pain modulation map to aberrant PAG connectivity

#### dl/lPAG and vlPAG showed distinct within-group connectivity patterns with other parts of the brain

As has previously reported in a smaller group of healthy controls^39^, the vlPAG and dl/lPAG showed significant differences in their whole brain connectivity. In brief, the dl/lPAG relative to vlPAG showed significantly higher connectivity with brain regions involved in threat detection, such as the thalamus, anterior cingulate cortex (ACC), and the dorsal and ventral attention networks. In comparison, the vlPAG showed significantly higher connectivity with sensory processing and memory processing regions. This pattern was overall consistent between the right and left columns, within each of the three groups (for details see supplementary section 9). This finding further confirms that human dl/l and vlPAG perform distinct functions related to threat processing.

#### High PDA group exhibits significant alterations in pain modulation circuitry

After correction for multiple comparisons, significant differences were observed in the comparison between the high PDA and HC groups. Significant differences in the high PDA group were localized to the right dl/lPAG, which showed increased negative connectivity with prefrontal and limbic regions, including the ACC, dmPFC (Dorsomedial Prefrontal Cortex), and nucleus accumbens (core and shell), compared to HC (Figure 4A,B). Additionally, a similar shift toward increased negative connectivity was observed for the left dl/lPAG, though limited to the left dmPFCa. For details, see Supplementary Table 8.

**Figure 4.**
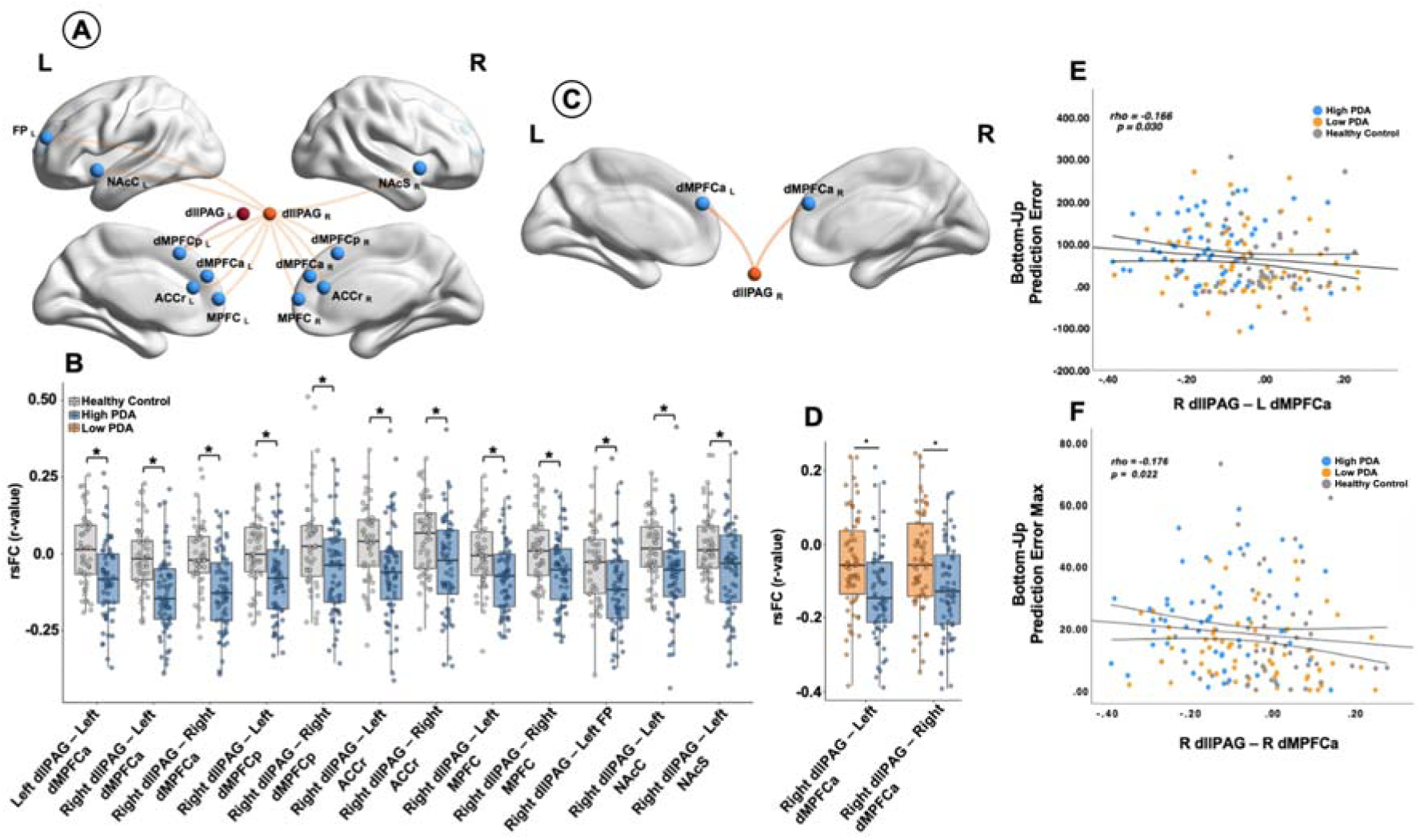
High PDA group showed significantly higher negative connectivity with other brain regions. ***A,*** Regions exhibiting significant group differences in rsFC with the right and left dl/lPAG between high PDA and HC (FDR corrected) are displayed. **B,** Box plots show the rsFC values (r) for significant regions. ***C,*** Among regions shown in (B) high and low PDA groups showed significant rsFC difference between the right dl/lPAG and dmPFCa (FDR corrected) ***D,*** Box plots illustrate rsFC values (r) for the high PDA (blue) and low PDA (orange) ***E,*** Shows the correlation between the right dl/lPAG–left dMPFCa contrast and bottom-up prediction error, while ***F*** Shows the correlation between the right dl/lPAG–right dMPFCa contrast and bottom-up prediction error max. Asterisks indicate significant group differences, p < 0.05. Data points are color-coded by group: blue for high PDA, orange for low PDA, and gray for HC. dMPFCa, dorsomedial prefrontal cortex (anterior division); dMPFCp, dorsomedial prefrontal cortex (posterior division); ACCr, anterior cingulate cortex (rostral); MPFC, medial prefrontal cortex; NAcC, nucleus accumbens core; nucleus accumbens shell; FP, frontal pole; dl/lPAG, dorsolateral/lateral periaqueductal gray.

After correction for multiple comparisons, the high and low PDA groups did not show a significant difference in PAG connectivity. We further checked whether the dl/lPAG connectivity patterns identified in the high PDA and HC contrast also show differences in connectivity between the high and low PDA groups. After FDR correction (q < 0.05), the high PDA group showed significantly more negative connectivity between the right dl/lPAG and both the left and right dmPFCa (*U* = 3383.00, *P* = 0.0009; *U* = 3482.00, *P* = 0.0045). (see supplementary Table 8 for full statistics and Figure 4C,D).

This pathway (dl/lPAG and dmPFCa) significantly predicted heightened responses to bottom-up prediction error (*ρ* = −0.166, *P* = 0.030; Figure 4E), and heightened bottom-up prediction error response max (*ρ* = −0.176, *P* = 0.022; Figure 4F). Additional correlations involving these connections are reported in Supplementary Figure 16.

Lastly, to assess whether demographic or motion-related factors influenced the observed rsFC differences between the high and low PDA groups, a multivariate general linear model (GLM) was conducted using the rank-transformed rsFC values from the seed-based analysis. The model included age (*P* = 0.902), sex (*P* = 0.776), framewise displacement (FD; *P* = 0.317), and DVARS (*P* = 0.585) as covariates. Thus, none of these variables had a significant effect on the main contrasts. In contrast, group membership showed a significant multivariate effect (*P* < 0.001), indicating that the rsFC differences were not attributable to age, sex, or motion-related confounds. We further confirmed the robustness of these findings by repeating the analysis with physiological noise correction (heart rate and respiration using RETROICOR), which replicated the reduced dl/lPAG–dmPFCa connectivity difference between the high and low PDA group (see Supplementary Section 11).

### Statistical modeling of high vs low PDA groups using logistic regression

To get an assessment of how well the significant behavioral, expectation task and PAG connectivity distinguishes between high and low PDA groups, we combined these metrics in a logistic regression model (see supplementary figure 18). The main goal of this analysis was to check whether combining the metrics gives a better accuracy than AUC (Area Under the Curve) based on the individual metrics and as a statistical comparator for subsequent machine learning analysis.

The overall model demonstrated a reasonable fit, as Indicated by a Nagelkerke *R²* = 0.247, explaining approximately 24.7% of the variance in group classification. Backward stepwise selection (Wald method) retained prediction error bias max and Right dl/lPAG – Left dMPFCa rsFC as the significant predictors (see supplementary Table 9). The model achieved a classification accuracy of 69.2%, with sensitivity at 68.4% and specificity at 70.0%, which was higher than the classification performance of any individual metric assessed in the ROC analysis.

### Stepwise machine learning models employing clinically feasible features differentiate PDA phenotypes

Three models were trained using a stepwise feature inclusion strategy to evaluate incremental performance gains and clinical feasibility. Table 2 summarizes cross-validation and test performance for logistic regression, SVM, and random forest classifiers across three feature sets in an order of increasingly lower clinical feasibility: (a) behavioral and clinical variables only, (b) behavioral plus schema-derived variables, and (c) all available features, including rsFC metrics. Since the objective of this study was to stratify low and high PDA, only the variables identified to distinguish the high PDA from the low PDA groups were used for ML.

**Table 2.**
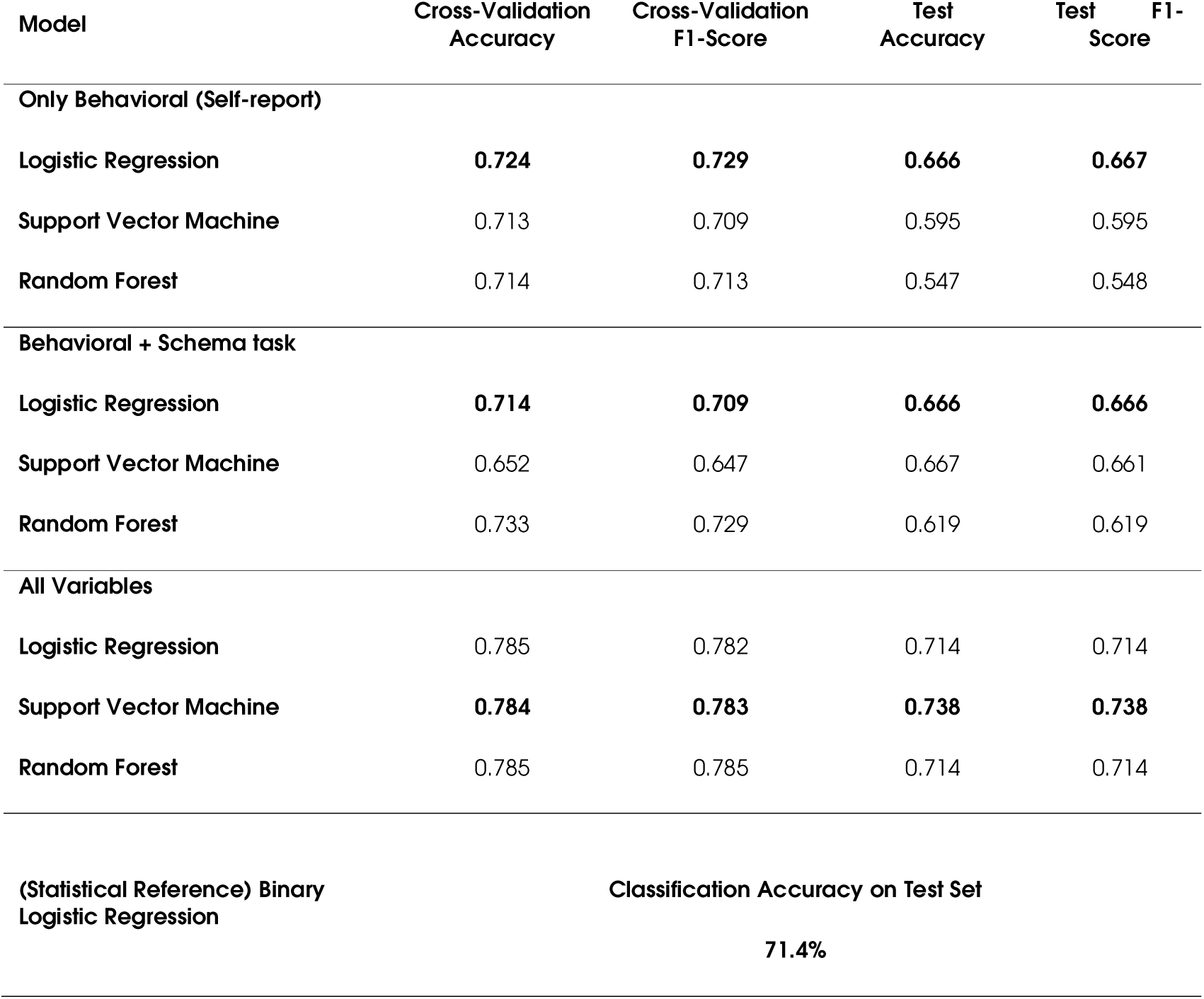
Stepwise Machine Learning Model Performance Metrics.

For the behavioral-only Model, 15 initial features were reduced to 10 based on statistical significance in distinguishing high vs. low PDA groups. Lasso regularization (*α* = 0.01) was then applied to penalize less informative predictors and avoid overfitting, and it retained all 10 features. Finally, correlation filtering (threshold *|r|* > 0.85) was used to remove highly collinear variables and reduce redundancy, leading to the exclusion of PCS and PVAQ subscales and yielding a final set of 6 features. Logistic regression showed the highest accuracy, followed by SVM and random forest (see Table 2)

For the Behavioral + Schema Model, two schema-derived features (BU prediction error (and max) responses), yielded 12 features, all of which were retained after Lasso regularization (optimal *α* = 0.01). Following correlation filtering, the final model included 8 predictors. The three models showed mild to modest gains relative to the behavioral-only model (See Table 2).

For the full model (Behavioral + Schema + dl/lPAG functional connectivity), 24 initial features were reduced to 14 through significance testing, all retained by Lasso (*α* = 0.001). Correlation filtering led to a final set of 10 predictors. As shown in Table 2, adding the neuroimaging predictors improved the classification performance to a maximum of 0.738 test accuracy with SVM.

As a statistics-based reference, we applied a standard binary logistic regression only to the test set as a statistical reference to contextualize the value of the machine learning approach. This approach yielded a test accuracy of 71.4%, which was lower than the performance of the best-performing machine learning classifier.

In terms of model interpretability, the best-performing SVM model underwent further assessment via confusion matrices and SHAP analysis. Confusion matrices (Figure 5b) showed accurate chronic pain severity differentiation in both training (*n* = 98) and test (*n* = 42) sets. On the independent test set, it correctly classified 17/23 high PDA and 14/19 low PDA. SHAP analysis (Figure 5A) identified the most influential predictors: McGill Sensory Score, MQS, and BPI areas of pain. Higher values of these features generally increased the likelihood of classification into the high PDA category.

**Figure 5.**
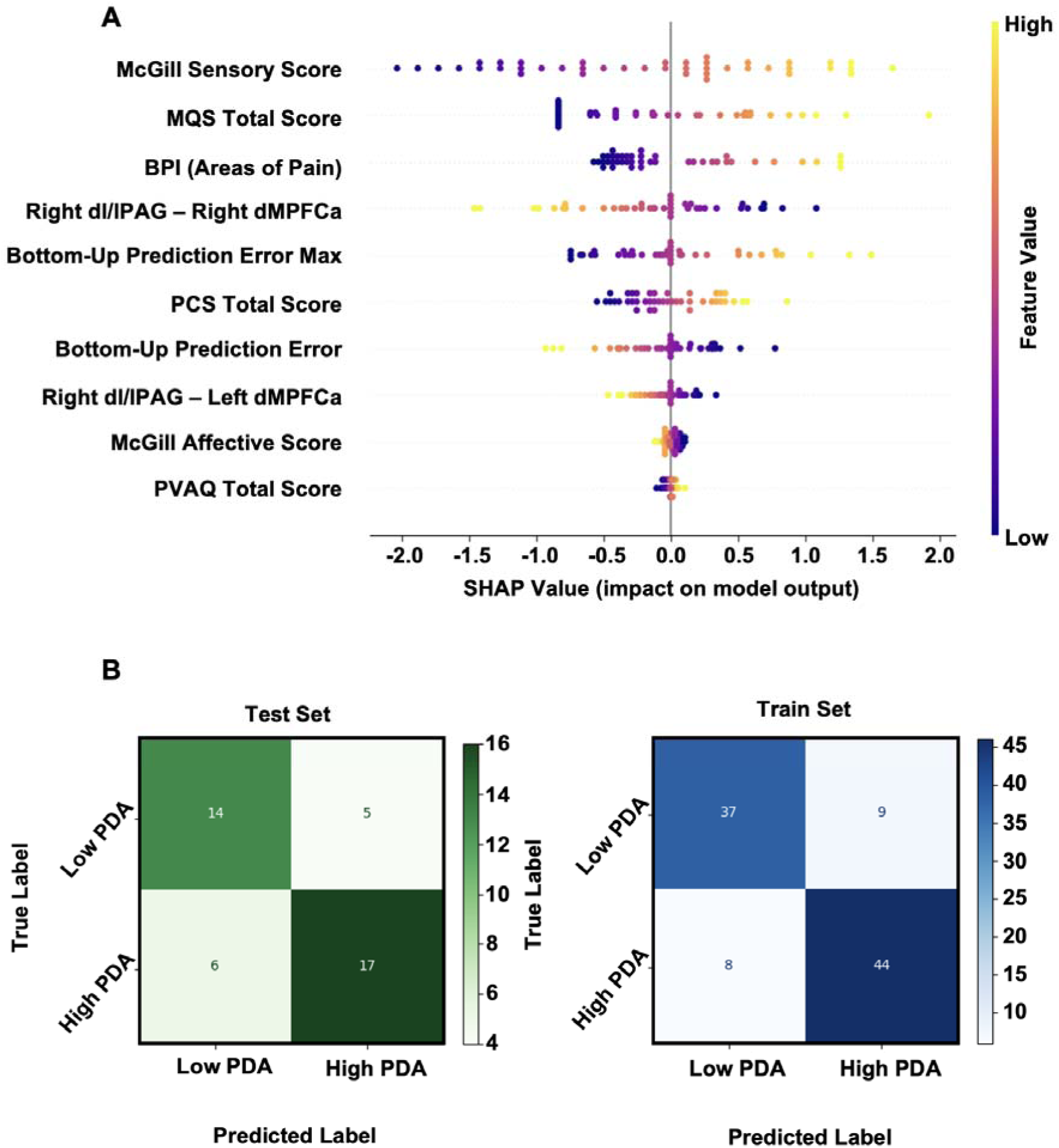
Interpretability and classification performance of the best-performing Support Vector Machine (SVM) model using the full feature set (behavioral, schema-related, and rsFC measures). ***A,*** SHAP (SHapley Additive Explanations) beeswarm plot displaying the relative contribution of each predictor to the SVM model’s classification output. Each point represents a participant, with color indicating the feature value (yellow = high, purple = low). Features are ordered by importance from top to bottom. Positive SHAP values correspond to higher likelihood of being classified into the high PDA group. Key contributing features include McGill Sensory Score, MQS Total Score, and the areas of pain (from BPI). *B,* Confusion matrices showing model performance on the independent test set (left; n = 42) and training set (right; n = 98). True labels are plotted on the y-axis and predicted labels on the x-axis. The model correctly classified 17 out of 23 individuals in the high PDA group and 14 out of 19 in the low PDA group on the test set indicating the generalizability and performance. BPI, Brief Pain Inventory; dl/lPAG; dorsolateral/lateral Periaqueductal gray; dMPFCa, anterior Dorsomedial Prefrontal Cortex; MQS, Medication Quantification Scale; PCS, Pain Catastrophizing Scale; PDA, Pain-Disability-Affect; PVAQ, Pain Vigilance and Awareness Questionnaire; SHAP, SHapley Additive exPlanations.

## Discussion

This study demonstrates that chronic pain subtypes, distinguished by differences in pain intensity, disability, and affective burden, are associated with distinct clinical, behavioral, and PAG connectivity profiles. These findings provide new insights into underlying mechanisms and identify key features with potential utility for distinguishing chronic pain phenotypes with differing clinical needs. Using core indicators of pain intensity, disability, and affective distress, we classified patients into high and low PDA groups, independent of diagnostic category. The high PDA group exhibited greater medication burden and significantly elevated fear-related responses, indicating that this group needs comprehensive treatment strategies. This group also showed impaired pain modulation in response to violated positive expectations and altered resting-state connectivity between the dl/lPAG and brain regions involved in cognitive appraisal. Machine learning models trained on selected clinical, behavioral, and neuroimaging features reliably distinguished individuals in the high and low PDA groups, supporting the utility of this framework for future stratification and targeted intervention efforts.

Recent nosological and conceptual developments in pain research have reframed chronic primary pain (CPP) as a disease entity in its own right, rather than a symptom of peripheral pathology. This shift, formalized by the ICD-11, reflects a growing consensus that conditions such as fibromyalgia and nonspecific chronic low back pain may arise from distinct biological mechanisms, including central sensitization, impaired descending modulation, and affective-cognitive dysregulation, rather than a single underlying cause.^65–67^ Despite this reframing, the neural mechanisms underlying CPP remain incompletely characterized, and current diagnostic categories do little to capture the observed heterogeneity across patients. Studies using quantitative sensory testing have identified divergent sensory profiles within and across diagnoses, but these mechanistic patterns have yet to be formally linked to symptom-based classification schemes.^68–70^ In comparison, multidimensional phenotyping frameworks that align with the biopsychosocial model of pain, conceptualizes pain as an emergent property of somatic, affective, and functional systems.^71,72^ Clinically, individuals with high PDA burden consistently show poorer outcomes across a range of pharmacological, rehabilitative, and psychological interventions, and are more likely to experience persistent disability.^73–76^ Although psychologically informed approaches such as cognitive behavioral therapy can benefit some high-burden patients, response is far from uniform, particularly among those with elevated catastrophizing or psychiatric comorbidity.^77,78^ These observations raise a critical and testable hypothesis: symptom-defined phenotypes such as high vs. low PDA may correspond to biologically meaningful subtypes that influence both prognosis and treatment responsiveness. Within this framework, dimensions such as pain intensity, emotional distress, and functional interference were proposed as clinically relevant axes for stratifying patients and tailoring interventions with symptom constellations that map more directly onto real-world clinical needs.^3,79,80^ Our findings combine symptom-based clustering with hypothesis-driven brain connectivity and hence advance stratified, neurobiology-based and evidence-driven care by using domain-informed, stepwise machine learning to reliably distinguish high- and low-PDA phenotypes. By starting with clinically feasible measures, then adding schema-test features, and finally incorporating brain connectivity guided by a specific hypothesis, we achieved a test accuracy of ∼74%. ML provides a powerful approach to classify, predict, and stratify chronic pain patients, offering a path toward mechanism-informed, precision treatments but robust, clinically feasible pipelines are still emerging.^7,81–91^ Our results therefore extend prior work by showing that clinically feasible and mechanistically informed features can sharpen chronic pain phenotyping.

Patients with high PDA exhibited exaggerated pain responses when positive expectations were violated, compared with both low PDA and healthy controls, indicating heightened cognitive sensitivity to prediction errors. Within the predictive-coding framework, such disruption may reflect aberrant precision-weighting, which is an imbalance in how the brain assigns confidence to sensory prediction errors versus prior expectations.^92,93^ Cognitive-affective vulnerabilities^94^ such as hypervigilance and catastrophizing likely amplify perceived threat and reactivity to unexpected pain, promoting over-engagement of sensory systems during prediction-error processing.^32,34,95,96^ In high PDA, amplified attention towards bottom-up input will result whenever the high weighting of catastrophic priors is violated by low threat cues, resulting in exaggerated pain. Persistent negative expectations may further reinforce maladaptive anticipation and response patterns, reflecting failures in adaptive recalibration and learning.^97,98^ Collectively, these findings delineate a precision- and prediction-error-based mechanism that differentiates chronic pain phenotypes from healthy controls and extends current understanding of disrupted pain-modulation processes in high-burden chronic pain.

Note that there was no significant difference in pain sensitivity or in responses to uncued heat stimuli, indicating that the observed effect reflects a cognitive sensitivity to violated expectations rather than a sensory sensitivity. Hence, high PDA patients may benefit most from interventions aimed at downregulating hypervigilance and strengthening the capacity for flexibly modulating evaluations of pain. Interventional therapy generally grounded in the fear-avoidance model, such as Pain Reprocessing Therapy,^99^ Cognitive-behavioral approaches^100^, and Acceptance and Commitment Therapy, have been shown to have promising effectiveness in reducing chronic pain by helping patients reinterpret pain as a non-dangerous brain signal, recalibrating threat appraisals, increasing psychological flexibility and reducing experiential avoidance, with growing evidence of benefit across chronic pain groups.^101,102^

Most human studies have treated the PAG as a unified structure, and when subregions are considered, the focus has typically been on the vlPAG as the primary mediator of pain modulation.^103–107^ In contrast, the dorsolateral/lateral PAG has received far less attention, despite evidence showing its distinct functional role in humans^108–110^ and animals.^111–114^ The current findings, together with previous work from our group^39^, indicates that dl/lPAG has an important role in threat processing. The altered connectivity between the dl/lPAG and dmPFC observed in the high PDA group aligns with emerging neuroimaging evidence suggesting that this subregion plays a distinct role in integrating aversive prediction errors with cognitive-affective regulators or processes during threat processing.^39,45^ These findings converge with previous evidence identifying the dorsal PAG as a key structure for aversive prediction error coding, responding more strongly when pain is worse than expected and communicating bidirectionally with medial prefrontal regions.^20^ In our study, high PDA individuals exhibited increased negative resting-state connectivity between the dl/lPAG and dmPFC, a pattern not observed in low PDA or control groups. Importantly, the same dl/lPAG–dmPFC connectivity pattern was significantly correlated with prediction error responses, suggesting a functional decoupling of regulatory input from prefrontal cortex to brainstem pain modulation circuits is directly linked to the abnormal expectation processing observed in high PDA. By identifying distinct patterns of pain modulation and dl/lPAG–dmPFC connectivity in high vs. low PDA groups, we offer mechanistic insight into individual differences in pain vulnerability.

Given the complexity of chronic pain, we incorporated validation steps to ensure the specificity of our results. The main connectivity results held even when smoothing was removed, after correction for heart rate and respiration artifacts. Moreover, group differences were unrelated to age, sex, or motion. Seed placement was verified, and a whole-brainstem region was tested as a control analysis and did not reproduce the dl/lPAG findings. While our machine learning approach prioritized interpretability and clinical relevance, future studies could benefit from validation using larger and blinded datasets to assess generalizability.

In summary, this study shows that chronic pain can be meaningfully subdivided into groups defined by pain intensity, disability, and affective distress. These subtypes align with distinct clinical, behavioral, and brain connectivity features that capture both emotional and sensory influences on pain regulation. High vs low PDA groups demonstrated different patterns of pain modulation and dl/lPAG–dmPFC connectivity, providing mechanistic insight into individual vulnerability to pain. These findings suggest that chronic pain subtypes can be identified using a mechanism-based approach informed by prior models and clinical observations.

## Data availability

Data will be made available in a public repository before the paper is published.

## Funding

The authors thank all participants who took part in the experiments. We would like to acknowledge our funding sources: the Natural Sciences and Research Engineering Council of Canada (NSERC) Discovery Grant, the Canada Research Chairs Program, the John R. Evans Leaders and Canada Innovation Funds (CFI-JELF), the Canadian Institute of Health Research (CIHR) Project Grant.

## Competing interests

The authors report no competing interests.

## Supplementary material

Supplementary material is available at *Brain* online.

## Notes

### Competing Interest Statement

The authors have declared no competing interest.

