## Supplementary material for "Pain-regulation circuitry as a predictor of chronic pain phenotypes": supplementary.pdf

### Supplementary Materials

#### 1. Flowchart of participants

Flowchart detailing participant inclusion and exclusion throughout the study. A total of 231 participants were initially recruited, including 159 individuals with chronic pain (CLBP = 109, FM = 50) and 72 healthy controls. Based on symptom clustering, 11 chronic pain participants (FM = 3, CLBP = 8) did not fit into either the high or low Pain–Disability–Affect (PDA) clusters and were excluded. This resulted in 75 participants in the high PDA cluster (FM = 34, CLBP = 41) and 73 in the low PDA cluster (FM = 13, CLBP = 60). Following initial age and sex matching, 20 participants were excluded (12 healthy controls and 8 CLBP participants in the low PDA cluster). An additional 8 participants were removed during resting-state fMRI quality control due to excessive motion, abnormal DVARS, or missing rs-fMRI data (5 healthy controls, 2 CLBP high PDA, and 1 CLBP low PDA). A final age and sex matching step led to the exclusion of 13 more participants (2 healthy controls, 9 high PDA [5 FM, 4 CLBP], and 2 low PDA [1 FM, 1 CLBP]). The final sample included 53 healthy controls, 64 participants in the high PDA cluster, and 62 in the low PDA cluster for inclusion in the rs-fMRI analyses.

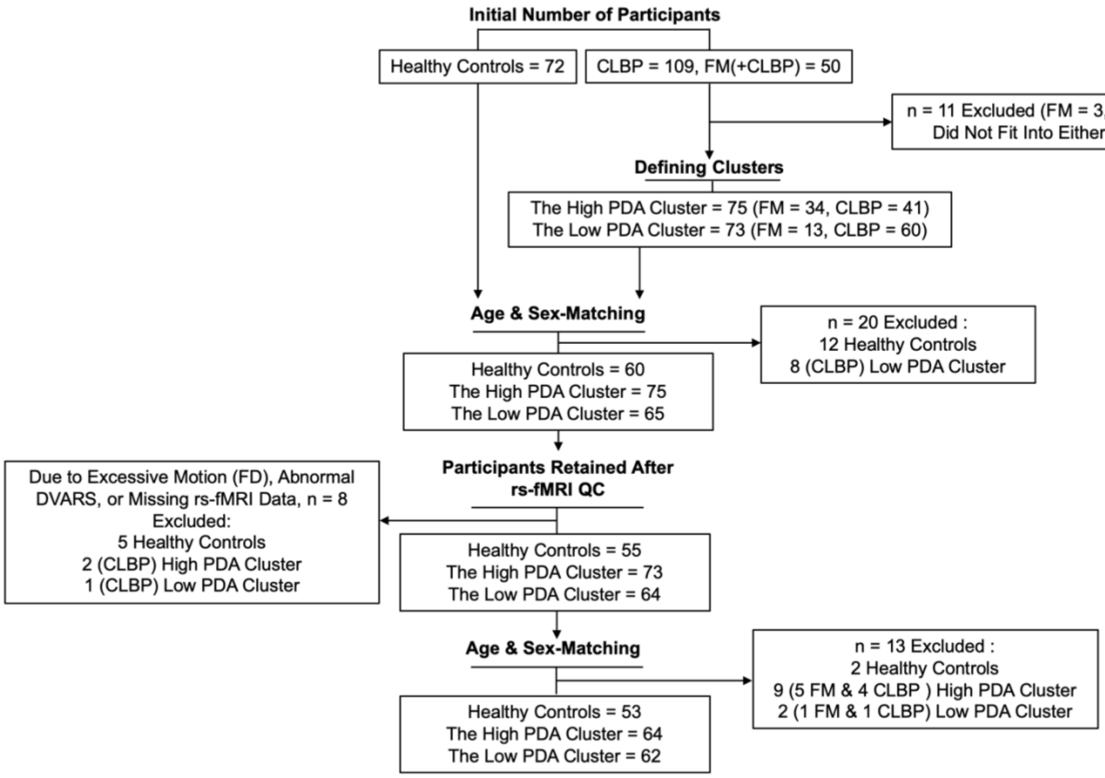

***Supplementary Figure 1.*** Participant inclusion and exclusion flowchart. CLBP, chronic low back pain; FM, fibromyalgia; PDA, Pain–Disability–Affect; FD, framewise displacement; DVARS, temporal derivative of root mean square variance across voxels; rs-fMRI, resting-state functional magnetic resonance imaging.

#### 2. Details on Principal Component Analysis and Clustering

PCA was performed to reduce data dimensionality across pain impact and affective distress measures. Suitability tests confirmed its applicability (KMO = 0.691; Bartlett's test: $\chi^2 = 354.90$, df = 10, $p < 0.001$). Two components with eigenvalues >1 were extracted, jointly explaining 83.04% of the variance. The first component (eigenvalue = 2.82, variance = 56.47%) had strong loadings on BDI, state anxiety, and trait anxiety, while the second component (eigenvalue = 1.33, variance = 26.57%) primarily reflected BPI pain severity and interference. The scree plot (Supplementary Figure 2) confirmed this two-component solution.

After Varimax rotation, the first component retained high loadings on BDI (0.886), state anxiety (0.884), and trait anxiety (0.939), isolating it as affective distress. The second component showed strong loadings on BPI pain severity (0.912) and pain interference (0.873), defining it as pain impact. This two-factor structure was retained for clustering analysis.

As a next step, hierarchical clustering, using Euclidean distance, was applied to determine the optimal number of clusters for k-means clustering. Using Ward's linkage method**,** a dendrogram (Supplementary Figure 3) revealed a clear bifurcation into two clusters, indicating that a two-group structure best captured the underlying heterogeneity in symptom presentation. This informed a subsequent K-means clustering analysis, which iteratively partitioned the data by minimizing within-cluster variance and maximizing separation between groups (see Supplementary Figure 4 for an overview of the clustering pipeline).

K-means clustering was performed with $k = 2$, successfully converging after eight iterations. The final cluster centers differentiated participants based on symptom severity, with one group exhibiting higher scores on affective distress (0.29143) and pain impact (0.57008), while the other displayed lower scores on affective distress (-0.47878) and pain impact (-0.93657). This separation was statistically confirmed, with ANOVA revealing significant differences between the two groups for both PCA components (Component 1: $F(1, 146) = 23.86$, $p < .001$, MS = 20.65; Component 2: $F(1, 146) = 169.71$, $p < .001$, MS = 79.02). The final cluster centers were separated by a Euclidean distance of 1.692, reflecting a distinct yet clinically relevant differentiation between the two groups. Despite the clear clustering structure, a subset of participants ($n = 11$; 3 FM, 8 CLBP) did not fit neatly within either severity-based group and were excluded from further analyses.

To further assess if the clustering approach effectively captured distinct symptom-based groups, the robustness of this two-cluster solution was determined using silhouette analysis. The high PDA group exhibited a mean silhouette score of 0.591, indicating strong within-cluster cohesion and clear separation, whereas the low PDA group had a slightly lower silhouette score of 0.537. The overall mean silhouette score of 0.567 suggests moderate clustering quality, with most participants aligning well within their respective groups (see supplementary figure 5).

A

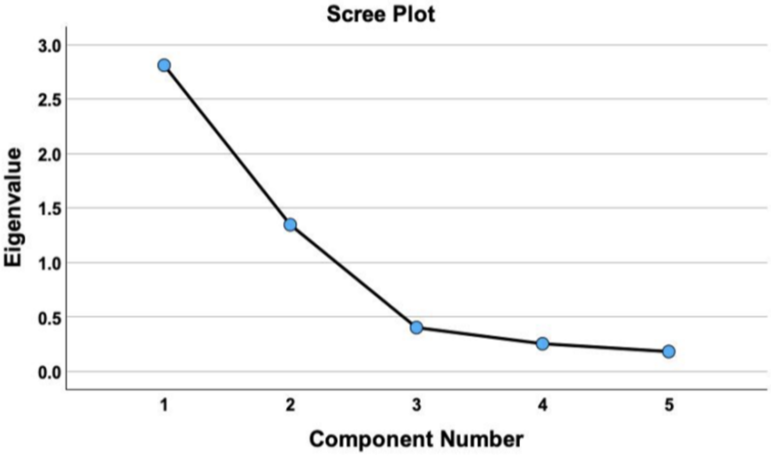

B

| | Component | |
|---|---|---|
| | 1 | 2 |
| BDI | 0.881 | -0.226 |
| State Anxiety | 0.850 | -0.282 |
| Trait Anxiety | 0.859 | -0.383 |
| BPI Pain Severity | 0.465 | 0.786 |
| BPI Pain Interference | 0.599 | 0.672 |

***Supplementary Figure 2.*** Principal Component Analysis (PCA) Scree Plot and Component Loadings Prior to Rotation. ***A,*** Scree plot illustrating the eigenvalues of each component, used to determine the number of components to retain. The plot shows a clear inflection point after the second component, indicating that two components account for the majority of variance, justifying their selection for further analysis. ***B,*** Component loading matrix prior to Varimax rotation, showing the initial associations of each variable with the two identified components. Component 1 exhibits high loadings on affective distress variables: BDI (0.881), State Anxiety (0.850), and Trait Anxiety (0.859), suggesting this component represents psychological distress. Component 2 shows stronger loadings on pain-related indices: BPI Pain Severity (0.786) and BPI Pain Interference (0.672), indicating a focus on pain impact.

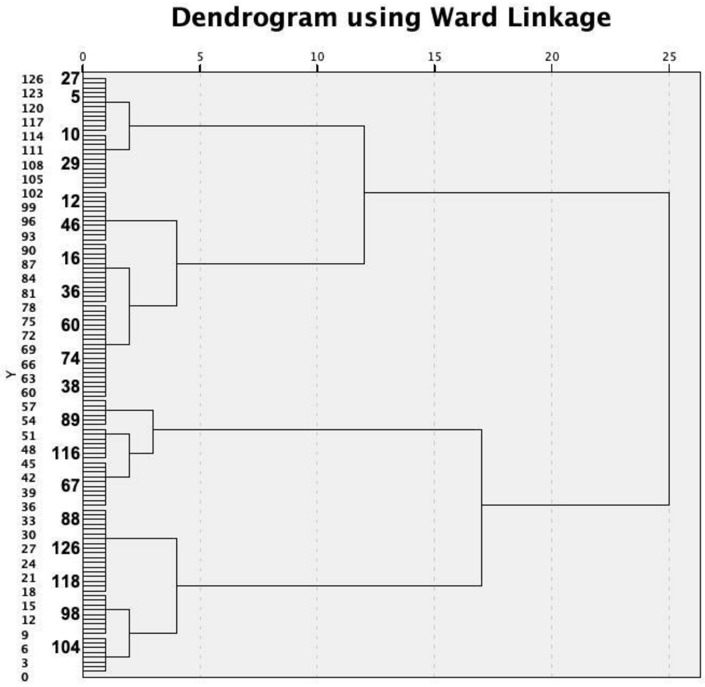

***Supplementary Figure 3.*** The dendrogram illustrates the hierarchical clustering of the dataset using Ward's linkage method, with distances measured along the y-axis. Each horizontal line represents the merging of clusters, with the height indicating the distance or dissimilarity at which clusters are joined. The most significant separation, or the largest jump in the dendrogram, occurs when the dataset is divided into two main clusters, as highlighted by the major bifurcation in the diagram. This division reflects the most distinct grouping within the data, suggesting the presence of two primary underlying structures. This informed the subsequent application of K-means clustering, where two clusters were defined as the optimal number for further analysis. The approach aligns with standard practices in cluster analysis, ensuring that the clustering process remains unbiased and data-driven.

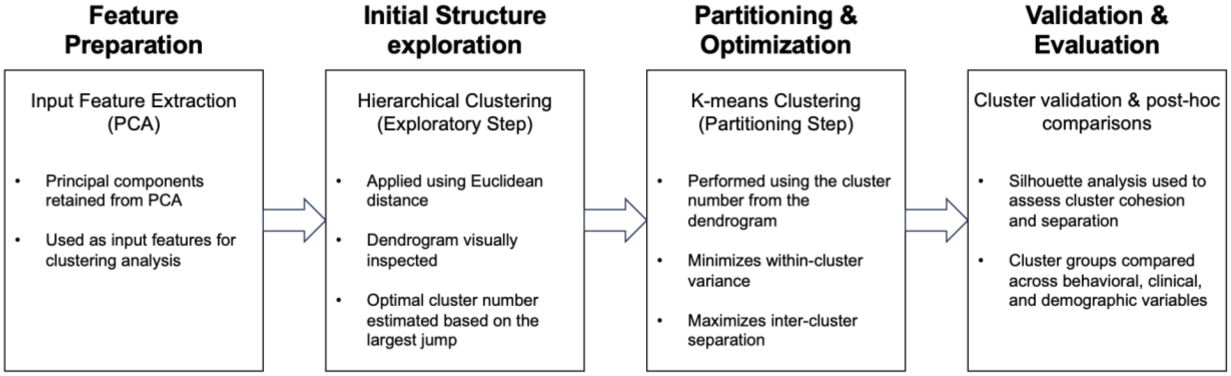

***Supplementary Figure 4.*** Schematic of the clustering pipeline used for symptom-based subgrouping. Principal components from pain, interference, and affective distress measures were entered into hierarchical and k-means clustering. Cluster validity was confirmed through silhouette analysis and post-hoc comparisons across behavioral, clinical, and demographic variables.

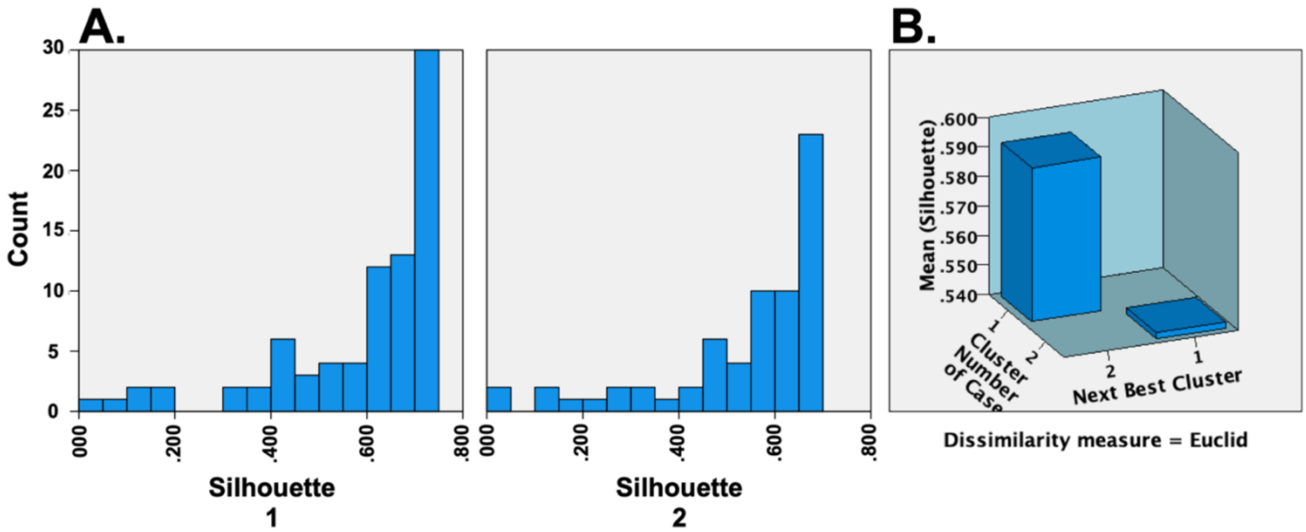

***Supplementary Figure 5.*** Silhouette Analysis for K-means Clustering Results. ***A,*** Distribution of silhouette scores by cluster. The histogram illustrates the distribution of silhouette values within each cluster, with Cluster 1 consisting of 75 cases and showing a mean silhouette score of 0.591 (range: 0.051 to 0.731). Cluster 2, which includes 73 cases, has a mean silhouette score of 0.537 (range: 0.024 to 0.682). Higher silhouette scores indicate better-defined clusters, with most cases in Cluster 1 and Cluster 2 exhibiting scores above 0.5, suggesting good separation between the clusters. ***B,*** Mean silhouette scores by cluster and next best cluster. The 3D bar plot compares the mean silhouette scores for each cluster against the next best cluster. The overall mean silhouette score across both clusters is 0.567, reinforcing the validity of the two-cluster solution. The next best cluster shows significantly lower silhouette scores, supporting the robustness of the current cluster assignments. Dissimilarity measure used: Euclidean distance.

### 3. MRI Data Acquisition and Preprocessing

#### 3.1. MRI data acquisition

Structural and functional MRI scans were collected using a 3T MRI scanner (Discovery MR750; General Electric Medical Systems, Waukesha, WI, USA) equipped with a 32-channel head coil (MR Instruments, Inc.; Minneapolis, MN, USA). The scanning sessions took place at the Halifax Infirmary Site, QEII Health Sciences Centre, Halifax, NS, Canada. To ensure participant comfort and minimize motion, foam padding was used to secure the head, and earplugs were provided to reduce the noise level from the scanner. Participants were also reminded to keep their heads still before each scan began. The resting-state scan lasted for 8 minutes.

The following parameters were used for acquiring T1-weighted structural brain images: field of view = 224 × 224 mm, in-plane resolution = 1 mm × 1 mm, slice thickness = 1.0 mm, TR/TE = 4.4/1.908 ms, and a flip angle of 9°. Functional MRI data were obtained using a BOLD (blood oxygenation level-dependent) sequence with a multiband EPI sequence protocol. The parameters for the fMRI acquisition were as follows: field of view = 216 × 216 mm, in-plane resolution = 3 mm × 3 mm, slice thickness = 3.0 mm, TR/TE = 950/30 ms, with a SENSE factor of 2 and an acceleration factor of 3. A total of 500 volumes were acquired during the resting-state scans. For distortion correction, reverse phase-encoded images were acquired to enable the application of FSL's top-up tool.

#### 3.2. Preprocessing of Resting-State Data

All functional datasets underwent correction for field map–based distortion. Preprocessing was carried out using AFNI (https://afni.nimh.nih.gov/afni) and FSL (https://www.fmrib.ox.ac.uk/fsl), with scripts provided by the 1,000 Functional Connectomes Project (https://www.nitrc.org/projects/fcon_1000). The preprocessing parameters were adapted from the parent study (3). The AFNI-based preprocessing steps included: (1) discarding the initial five volumes of each EPI run to allow for magnetization stabilization, (2) applying rigid-body motion correction by aligning each volume to the mean image via Fourier interpolation, (3) performing skull stripping, and (4) generating a representative eighth volume for registration purposes. FSL preprocessing included: (5) spatial smoothing with a 6 mm full-width at half-maximum Gaussian kernel, (6) grand-mean intensity normalization, (7) bandpass temporal filtering between 0.005 and 0.3 Hz, (8) removal of linear and quadratic trends, and (9) regression of nine nuisance signals (global signal, cerebrospinal fluid, white matter, and six motion parameters capturing head translation and rotation) in the native functional space.

Six motion parameters, corresponding to rotational movement and cardinal displacement, were generated during the FSL-based motion correction step in native functional space. Additionally, two nuisance time courses were calculated for white matter and ventricles using masks obtained from the image segmentation of the participant's T1-weighted data, applying a tissue-type probability threshold of 80%.

For standard space analysis, FLIRT was used to register the functional images to the MNI152 standard template (5). This process included: (a) registering the native-space structural image to the MNI152 2-mm template using a 12 degrees of freedom (df) linear affine transformation, (b) registering the native-space functional image to the high-resolution structural image using a 6 df linear transformation, and (c) computing the native-functional-to-standard-structural warps by concatenating the matrices computed in steps (a) and (b).

For data quality verification, maximum framewise displacement (FD) and DVARS (difference in signal intensity between successive volumes) were calculated to assess and exclude participants with excessive motion. The image quality criteria set a cutoff at a maximum FD above 3 mm or DVARS outliers detected in more than 10% of the acquired data (6). Based on these thresholds, four participants, two healthy controls and two chronic low back pain (CLBP) participants, were excluded from further analysis. Specifically, one HC and one CLBP participant exhibited excessive head motion, with maximum FD values of 6.71 mm and 4.44 mm, respectively. Additionally, one HC and one CLBP participant surpassed the DVARS threshold, with outliers detected in 11.74% and 18.01% of volumes, respectively. Following these exclusions, the highest remaining FD value within the included dataset was 2.62 mm, and the highest percentage of DVARS outliers was 9.91%, both within the acceptable range. To further control for residual motion artifacts, maximum FD and maximum DVARS were included as nuisance covariates in subsequent analyses correlating functional and behavioral metrics.

All main fMRI analyses were also repeated using data that was preprocessed without smoothing the intensities to confirm that the differences in dl/l and vlPAG results were not confounded due to smoothing (supplementary figure 6). Additionally, all main fMRI analyses were also repeated using a whole-brainstem ROI as a control region to confirm that observed PAG connectivity differences were not attributable to general connectivity within the brainstem (supplementary figure 7).

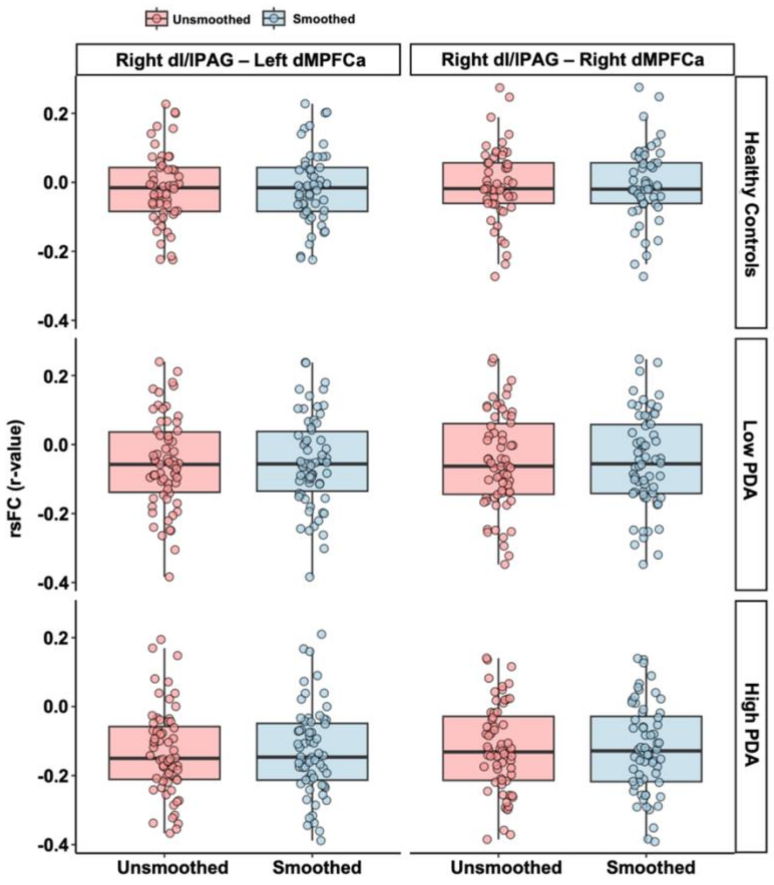

***Supplementary Figure 6.*** Control analysis using unsmoothed fMRI data confirms that main rsFC findings are unrelated to this reprocessing step. To assess the influence of spatial smoothing, all primary resting-state functional connectivity (rsFC) analyses were repeated using unsmoothed data. To illustrate, the figure shows rsFC (r-values) between the right dl/lPAG and bilateral dMPFCa, two key connections that significantly differentiated the high PDA group from both healthy controls and the low PDA group in the main analysis. Results are plotted separately for smoothed (blue) and unsmoothed (red) preprocessing pipelines across all three groups. The consistent pattern of effects across smoothing conditions confirms that these group differences are not attributable to preprocessing artifacts. rsFC differences between healthy controls and the high PDA group, were reproducible between smoothed and unsmoothed rsFC values, (data not shown).

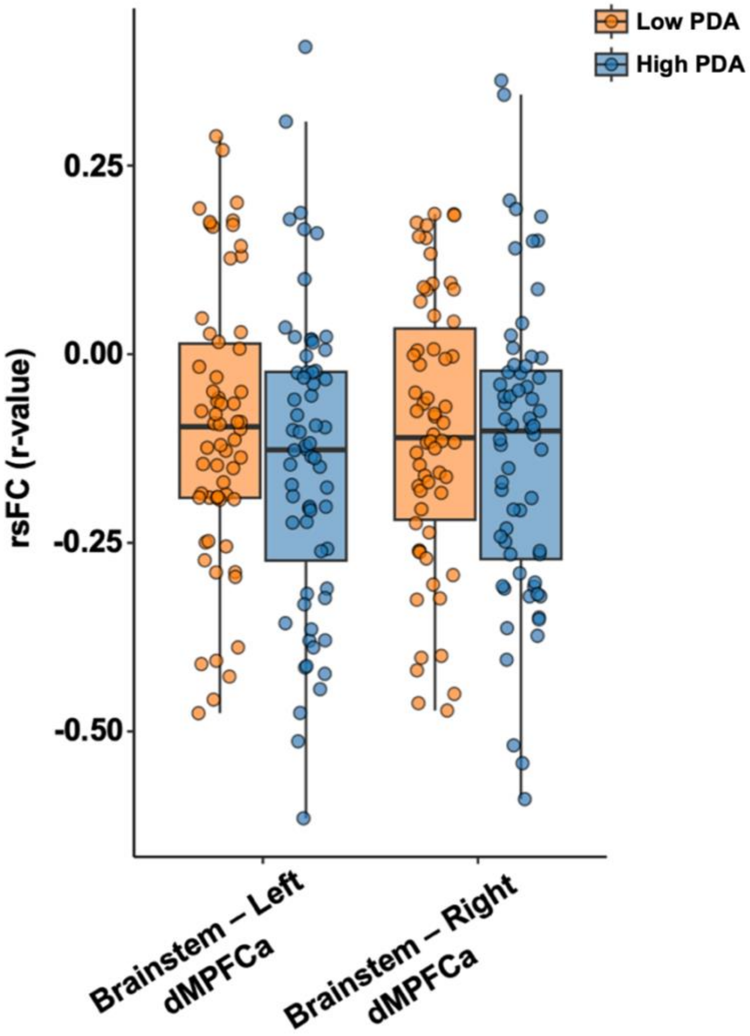

***Supplementary Figure 7.*** Control analysis using a whole-brainstem seed supports the anatomical specificity of dl/lPAG connectivity effects. To verify that group differences in PAG-based connectivity were not driven by generalized brainstem signals, a control analysis was conducted using the entire brainstem as the seed region. Resting-state functional connectivity (rsFC; Fisher's z-transformed r-values) with left and right dMPFCa is shown for the high and low PDA groups. Unlike the original analyses using the right dl/lPAG as a seed, no significant group differences were observed. This null finding suggests that the observed effects were not attributable to nonspecific or noise-related brainstem-wide connectivity patterns.

#### 4. PAG Seed Parcellation

The parcellation of the PAG was performed using a two-fold approach. Initially, focal PAG seed coordinates were selected based on previously published studies [1,2]. The primary coordinates for the dorsolateral/lateral (dl/lPAG) and ventrolateral PAG (vlPAG) seeds were x = ±4, y = −31, z = −8, and x = ±3, y = −32, z = −12, respectively. Masks for these PAG seeds were created under the following conditions: (1) a 3D voxel, (2) a selection size of 1, (3) a search radius of 1 [3], and (4) in Montreal Neurological Institute (MNI) 2mm$^3$ standard space. To further refine the seed placements, these masks were overlapped with an atlas previously created by [4], which was based on voxel-wise diffusion MRI and probabilistic tractography. Initially, data were analyzed separately for the left and right PAG columns. The spatial distribution of the PAG seed masks is illustrated in the supplementary figure 8. Additionally, registrations were visually checked to ensure that the PAG masks were registered properly within the brain stem gray area (Supplementary figure 9).

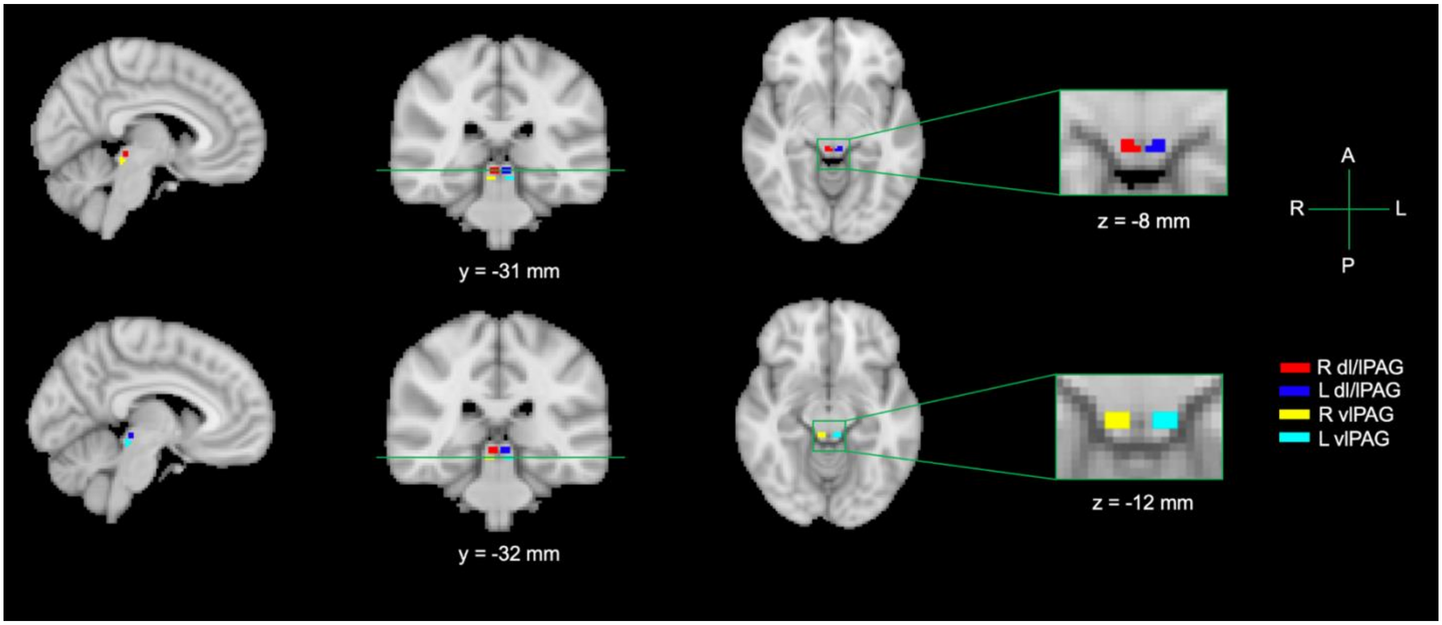

***Supplementary Figure 8.*** Parcellation of the periaqueductal gray (PAG) into dorsolateral/lateral (dl/lPAG) and ventrolateral (vlPAG) subdivisions. The dl/lPAG (red: right, blue: left) and vlPAG (yellow: right, cyan: left) seed masks were defined based on previously published MNI coordinates (dl/lPAG: x = ±4, y = −31, z = −8; vlPAG: x = ±3, y = −32, z = −12) and further refined using a probabilistic atlas. Coronal and axial slices illustrate the seed mask locations within the PAG in the MNI space. R = right; L = left; A = anterior; P = posterior.For more details, see Wang et al.[5] and Veinot et al.[6])

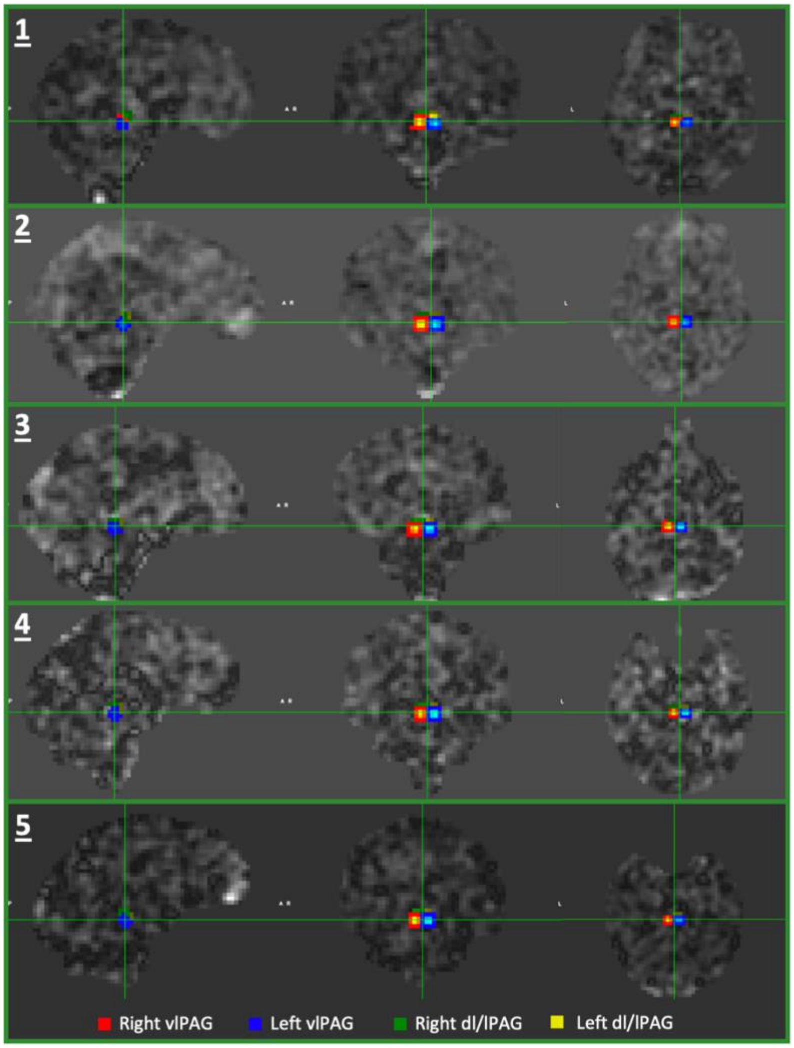

***Supplementary Figure 9***. Verification of PAG seed mask alignment in native functional space across representative participants selected randomly. Overlays of dl/lPAG (green = right, yellow = left) and vlPAG (red = right, blue = left) seed masks are shown on individual participants' unsmoothed EPI volumes across sagittal, coronal, and axial planes. Visual inspection confirmed consistent and anatomically appropriate placement of all seeds within the periaqueductal gray across subjects. This quality control step was performed to validate the transformation of MNI-space seed masks into native functional space and to ensure reliable localization of PAG subregions for downstream connectivity analyses.

***Supplementary Table 1.*** Parcellation scheme and MNI coordinates of regions of interest (ROIs). The Harvard–Oxford cortical and subcortical atlas served as the anatomical foundation for ROI definition, with all bilateral regions separated into distinct left and right hemispheric ROIs. Regions superior to the paramedial sulcus were designated as dorsal medial prefrontal cortex (dMPFC) and further subdivided into anterior (dMPFCa) and posterior (dMPFCp) subregions. The region occupying the superior rostral sulcus below the paramedial sulcus was demarcated as medial prefrontal cortex (MPFC), and the inferior rostral sulcus was labeled ventral medial prefrontal cortex (vMPFC). The frontal pole (FP) was subdivided into a superior FP and an inferior orbitofrontal pole region, approximately above and below the frontomarginal sulcus, respectively. The single cingulate ROI was replaced by the seven-region parcellation of Beckmann et al.[7], and the insula was subdivided into posterior, middle, dorsal anterior, and ventral anterior subregions following Hashmi et al.[8] To enhance anatomical specificity in ventral striatal and amygdalar regions, the single nucleus accumbens (NAc) ROI was replaced by distinct core and shell subregions defined according to the connectivity-based parcellation of Cartmell et al.[9], and the whole NAc region was excluded to avoid redundancy; the selected coordinates are consistent with those used in studies applying similar core–shell distinctions.[10–12] Likewise, the whole amygdala ROI was excluded and replaced by basolateral (BLA) and central (CeA) subregions derived from the connectivity-based parcellation of Bzdok et al.[13], in line with subsequent functional imaging work using similar subdivisions.[14]

| NO. | REGIONS | ABBREVIATIONS | MNI X | MNI Y | MNI Z |
|---|---|---|---|---|---|
| 1 | Caudal anterior cingulate left | ACCc_L | -4 | 40 | -2 |
| 2 | Caudal anterior cingulate right | ACCc_R | 4 | 40 | -2 |
| 3 | Mid anterior cingulate left | ACCm_L | -6 | -2 | 42 |
| 4 | Mid anterior cingulate right | ACCm_R | 6 | -2 | 42 |
| 5 | Rostral anterior cingulate left | ACCr_L | -4 | 38 | 18 |
| 6 | Rostral anterior cingulate mid posterior left | ACCrm_L | -6 | 18 | 34 |
| 7 | Rostral anterior cingulate mid posterior right | ACCrm_R | 6 | 18 | 34 |
| 8 | Rostral anterior cingulate posterior left | ACCrp_L | -4 | 22 | 20 |
| 9 | Rostral anterior cingulate posterior right | ACCrp_R | 4 | 22 | 20 |
| 10 | Rostral anterior cingulate right | ACCr_R | 4 | 38 | 18 |
| 11 | Subgenual anterior cingulate left | ACCsg_L | -4 | 16 | -14 |
| 12 | Subgenual anterior cingulate right | ACCsg_R | 4 | 16 | -14 |
| 13 | Angular gyrus left | Ang_L | -54 | -56 | 26 |
| 14 | Angular gyrus right | Ang_R | 54 | -56 | 26 |
| 15 | Brain stem | BrStem | 0 | -26 | -28 |
| 16 | Caudate left | Caud_L | -12 | 14 | 8 |
| 17 | Caudate right | Caud_R | 12 | 14 | 8 |
| 18 | Cingulate gyrus, posterior division left | Cingp_L | -4 | -38 | 32 |
| 19 | Cingulate gyrus, posterior division right | Cingp_R | 4 | -38 | 32 |
| 20 | Central opercular cortex left | Cop_L | -48 | -4 | 8 |
| 21 | Central opercular cortex right | Cop_R | 48 | -4 | 8 |
| 22 | Cuneal cortex left | Cun_L | -4 | -82 | 30 |
| 23 | Cuneal cortex right | Cun_R | 4 | -82 | 30 |
| 24 | Dorsal anterior insula left | dINSa_L | -32 | 20 | 0 |
| 25 | Dorsal anterior insula right | dINSa_R | 32 | 20 | 0 |
| 26 | Dorsal medial prefrontal cortex, anterior division left | dMPFCa_L | -4 | 50 | 28 |
| 27 | Dorsal medial prefrontal cortex, anterior division right | dMPFCa_R | 4 | 50 | 28 |
| 28 | Dorsal medial prefrontal cortex, posterior division left | dMPFCp_L | -4 | 26 | 48 |

| | | | | | |
|---|---|---|---|---|---|
| **29** | Dorsal medial prefrontal cortex, posterior division right | dMPFCp_R | 4 | 26 | 48 |
| **30** | Frontal orbital cortex left | FO_L | -40 | 30 | -14 |
| **31** | Frontal operculum cortex left | Fop_L | -40 | 20 | 4 |
| **32** | Frontal operculum cortex right | Fop_R | 40 | 20 | 4 |
| **33** | Frontal orbital cortex right | FO_R | 40 | 30 | -14 |
| **34** | Frontal pole left | FP_L | -30 | 54 | 20 |
| **35** | Frontal pole right | FP_R | 30 | 54 | 20 |
| **36** | Globus pallidus left | GP_L | -16 | -2 | -2 |
| **37** | Globus pallidus right | GP_R | 16 | -2 | -2 |
| **38** | Heschls gyrus (includes H1 and H2) left | He_L | -48 | -18 | 6 |
| **39** | Heschls gyrus (includes H1 and H2) right | He_R | 48 | -18 | 6 |
| **40** | Hippocampus left | Hipp_L | -28 | -22 | -16 |
| **41** | Hippocampus right | Hipp_R | 28 | -22 | -16 |
| **42** | Intracalcarine cortex left | IC_L | -6 | -74 | 12 |
| **43** | Intracalcarine cortex right | IC_R | 6 | -74 | 12 |
| **44** | Inferior frontal gyrus, pars opercularis left | IFGpo_L | -54 | 14 | 16 |
| **45** | Inferior frontal gyrus, pars opercularis right | IFGpo_R | 54 | 14 | 16 |
| **46** | Inferior frontal gyrus, pars triangularis left | IFGpt_L | -50 | 30 | 16 |
| **47** | Inferior frontal gyrus, pars triangularis right | IFGpt_R | 50 | 30 | 16 |
| **48** | Middle insula left | INSm_L | -40 | -2 | -2 |
| **49** | Middle insula right | INSm_R | 40 | -2 | -2 |
| **50** | Posterior insula left | INSp_L | -38 | -14 | 8 |
| **51** | Posterior insula right | INSp_R | 38 | -14 | 8 |
| **52** | Inferior temporal gyrus, anterior division left | ITGa_L | -50 | -6 | -40 |
| **53** | Inferior temporal gyrus, anterior division right | ITGa_R | 50 | -6 | -40 |
| **54** | Inferior temporal gyrus, posterior division left | ITGp_L | -56 | -32 | -24 |
| **55** | Inferior temporal gyrus, posterior division right | ITGp_R | 56 | -32 | -24 |
| **56** | Inferior temporal gyrus, temporooccipital part left | ITGtp_L | -56 | -54 | -18 |
| **57** | Inferior temporal gyrus, temporooccipital part right | ITGtp_R | 56 | -54 | -18 |
| **58** | Lingual gyrus left | Ling_L | -10 | -68 | -2 |
| **59** | Lingual gyrus right | Ling_R | 10 | -68 | -2 |
| **60** | Lateral occipital cortex, inferior division left | LOcci_L | -48 | -78 | -2 |
| **61** | Lateral occipital cortex, inferior division right | LOcci_R | 48 | -78 | -2 |
| **62** | Lateral occipital cortex, superior division left | LOccs_L | -40 | -78 | 34 |
| **63** | Lateral occipital cortex, superior division right | LOccs_R | 40 | -78 | 34 |
| **64** | Middle frontal gyrus left | MFG_L | -40 | 20 | 44 |
| **65** | Middle frontal gyrus right | MFG_R | 40 | 20 | 44 |
| **66** | Medial prefrontal cortex left | MPFC_L | -6 | 60 | 8 |
| **67** | Medial prefrontal cortex right | MPFC_R | 6 | 60 | 8 |
| **68** | Middle temporal gyrus, anterior division left | MTGa_L | -58 | -2 | -22 |
| **69** | Middle temporal gyrus, anterior division right | MTGa_R | 58 | -2 | -22 |

| | | | | | |
|---|---|---|---|---|---|
| **70** | Middle temporal gyrus, posterior division left | MTGp_L | -62 | -22 | -18 |
| **71** | Middle temporal gyrus, posterior division right | MTGp_R | 62 | -22 | -18 |
| **72** | Middle temporal gyrus, temporooccipital part left | MTGto_L | -60 | -52 | 0 |
| **73** | Middle temporal gyrus, temporooccipital part right | MTGto_R | 60 | -52 | 0 |
| **74** | Occipital fusiform gyrus left | OccFG_L | -28 | -76 | -14 |
| **75** | Occipital fusiform gyrus right | OccFG_R | 28 | -76 | -14 |
| **76** | Occipital pole left | OccP_L | -8 | -100 | 6 |
| **77** | Occipital pole right | OccP_R | 8 | -100 | 6 |
| **78** | Orbito frontal pole left | OFP_L | -32 | 58 | -6 |
| **79** | Orbito frontal pole right | OFP_R | 32 | 58 | -6 |
| **80** | Precuneous cortex left | pCun_L | -4 | -64 | 38 |
| **81** | Precuneous cortex right | pCun_R | 4 | -64 | 38 |
| **82** | Parahippocampal gyrus, anterior division left | pHippa_L | -24 | -6 | -34 |
| **83** | Parahippocampal gyrus, anterior division right | pHippa_R | 24 | -6 | -34 |
| **84** | Parahippocampal gyrus, posterior division left | pHippp_L | -24 | -32 | -18 |
| **85** | Parahippocampal gyrus, posterior division right | pHippp_R | 24 | -32 | -18 |
| **86** | Planum polare left | PIP_L | -48 | -4 | -6 |
| **87** | Planum polare right | PIP_R | 48 | -4 | -6 |
| **88** | Planum temporale left | PIT_L | -60 | -22 | 8 |
| **89** | Planum temporale right | PIT_R | 60 | -22 | 8 |
| **90** | Parietal operculum cortex left | Pop_L | -48 | -32 | 20 |
| **91** | Parietal operculum cortex right | Pop_R | 48 | -32 | 20 |
| **92** | Postcentral gyrus left | PostC_L | -54 | -20 | 46 |
| **93** | Postcentral gyrus right | PostC_R | 54 | -20 | 46 |
| **94** | Precentral gyrus left | PreC_L | -44 | -8 | 52 |
| **95** | Precentral gyrus right | PreC_R | 44 | -8 | 52 |
| **96** | Putamen left | Put_L | -30 | -4 | 0 |
| **97** | Putamen right | Put_R | 30 | -4 | 0 |
| **98** | Supracalcarine cortex left | Sc_L | -2 | -84 | 12 |
| **99** | Supracalcarine cortex right | Sc_R | 2 | -84 | 12 |
| **100** | Superior frontal gyrus left | SFG_L | -22 | 22 | 54 |
| **101** | Superior frontal gyrus right | SFG_R | 22 | 22 | 54 |
| **102** | Supplementary motor area left | SMA_L | -4 | -2 | 58 |
| **103** | Supplementary motor area right | SMA_R | 4 | -2 | 58 |
| **104** | Supra marginal gyrus left | SMGa_L | -58 | -32 | 40 |
| **105** | Supra marginal gyrus right | SMGa_R | 58 | -32 | 40 |
| **106** | Supramarginal gyrus, posterior division left | SMGp_L | -60 | -48 | 32 |
| **107** | Supramarginal gyrus, posterior division right | SMGp_R | 60 | -48 | 32 |
| **108** | Superior parietal lobule left | SPL_L | -32 | -50 | 60 |
| **109** | Superior parietal lobule right | SPL_R | 32 | -50 | 60 |
| **110** | Superior temporal gyrus, anterior division left | STGa_L | -58 | -4 | -6 |

| | | | | | |
|---|---|---|---|---|---|
| **111** | Superior temporal gyrus, anterior division right | STGa_R | 58 | -4 | -6 |
| **112** | Superior temporal gyrus, posterior division left | STGp_L | -66 | -26 | 6 |
| **113** | Superior temporal gyrus, posterior division right | STGp_R | 66 | -26 | 6 |
| **114** | Temporal fusiform cortex, anterior division left | TFCa_L | -32 | -6 | -42 |
| **115** | Temporal fusiform cortex, anterior division right | TFCa_R | 32 | -6 | -42 |
| **116** | Temporal fusiform cortex, posterior division left | TFCp_L | -36 | -16 | -32 |
| **117** | Temporal fusiform cortex, posterior division right | TFCp_R | 36 | -16 | -32 |
| **118** | Thalamus left | Thal_L | -10 | -18 | 8 |
| **119** | Thalamus right | Thal_R | 10 | -18 | 8 |
| **120** | Temporal occipital fusiform cortex left | TOF_L | -34 | -54 | -16 |
| **121** | Temporal occipital fusiform cortex right | TOF_R | 34 | -54 | -16 |
| **122** | Temporal pole left | TP_L | -40 | 16 | -30 |
| **123** | Temporal pole right | TP_R | 40 | 16 | -30 |
| **124** | Ventral anterior insula left | vINSa_L | -36 | 10 | -14 |
| **125** | Ventral anterior insula right | vINSa_R | 36 | 10 | -14 |
| **126** | Ventral medial prefrontal cortex left | vMPFC_L | -4 | 50 | -20 |
| **127** | Ventral medial prefrontal cortex right | vMPFC_R | 4 | 50 | -20 |
| **128** | Nucleus Accumbens Core left | NAcC_L | -10 | -14 | -6 |
| **129** | Nucleus Accumbens Core right | NAcC_R | 10 | 14 | -6 |
| **130** | Nucleus Accumbens Shell left | NAcS_L | -8 | 10 | -9 |
| **131** | Nucleus Accumbens Shell right | NAcS_R | 8 | 10 | -9 |
| **132** | Basolateral Amygdala left | BLA_L | -22 | -10 | -28 |
| **133** | Basolateral Amygdala right | BLA_R | 22 | -10 | -28 |
| **134** | Central Nucleus of Amygdala left | CeA_L | -20 | -6 | -14 |
| **135** | Central Nucleus of Amygdala right | CeA_R | 20 | -6 | -14 |

#### 5. Stepwise Machine Learning Models comparing features based on feasibility

To evaluate whether domain knowledge–driven feature extraction can support classification of chronic pain severity, we implemented a stepwise supervised machine learning strategy (see Supplementary Figure 10).

Three models were trained using progressively inclusive predictor sets, reflecting the feasibility of clinical data acquisition. Features were selected based on their statistical significance and clinical relevance, informed by hypothesis testing and expert judgment. The models were trained on: (1) behavioral and clinical variables alone; (2) those plus schema-related predictors from the pain modulation task; and (3) all of the above plus resting-state functional connectivity (rsFC) metrics from neuroimaging.

Standard preprocessing steps were applied across all datasets: non-numeric placeholders were converted to missing values, all variables were cast to numeric formats, and missing data were imputed using feature-wise means to avoid introducing bias.

Each model underwent independent feature selection within its own predictor set to ensure fair comparison across models with varying input complexity. To reduce dimensionality, Lasso regression with five-fold cross-validation was applied to each predictor group. This L1-regularized method shrinks less informative features to zero, promoting model sparsity. A predefined alpha grid (0.001, 0.01, 0.1, 1, 10) was tested to find the optimal regularization strength, and only features with non-zero coefficients were retained.

To further eliminate redundancy, we computed pairwise correlations among the retained features. Highly correlated pairs ($|r| > 0.85$) were reviewed, and in cases of conceptual overlap (e.g. subscale vs. total scores) the more interpretable or comprehensive measure was kept, preserving parsimony and clinical clarity.[15–18]

The data were then split into training and test sets using a stratified 70–30 split, maintaining balanced representation of severity classes. Within the training set, a five-fold cross-validation approach was used to optimize model performance and mitigate overfitting. For each feature set, three classifiers (logistic regression, random forest, and SVM) were trained and tuned via grid search across relevant hyperparameter spaces. Optimal configurations were chosen based on cross-validation accuracy and weighted F1-score.

The final performance was evaluated on the independent test set. Accuracy and weighted F1-score were used to assess overall performance, while confusion matrices visualized class-level prediction outcomes. For the top-performing model, SHapley Additive exPlanations (SHAP) were used to interpret feature contributions, and a beeswarm plot illustrated the direction and importance of each predictor's influence on individual predictions.

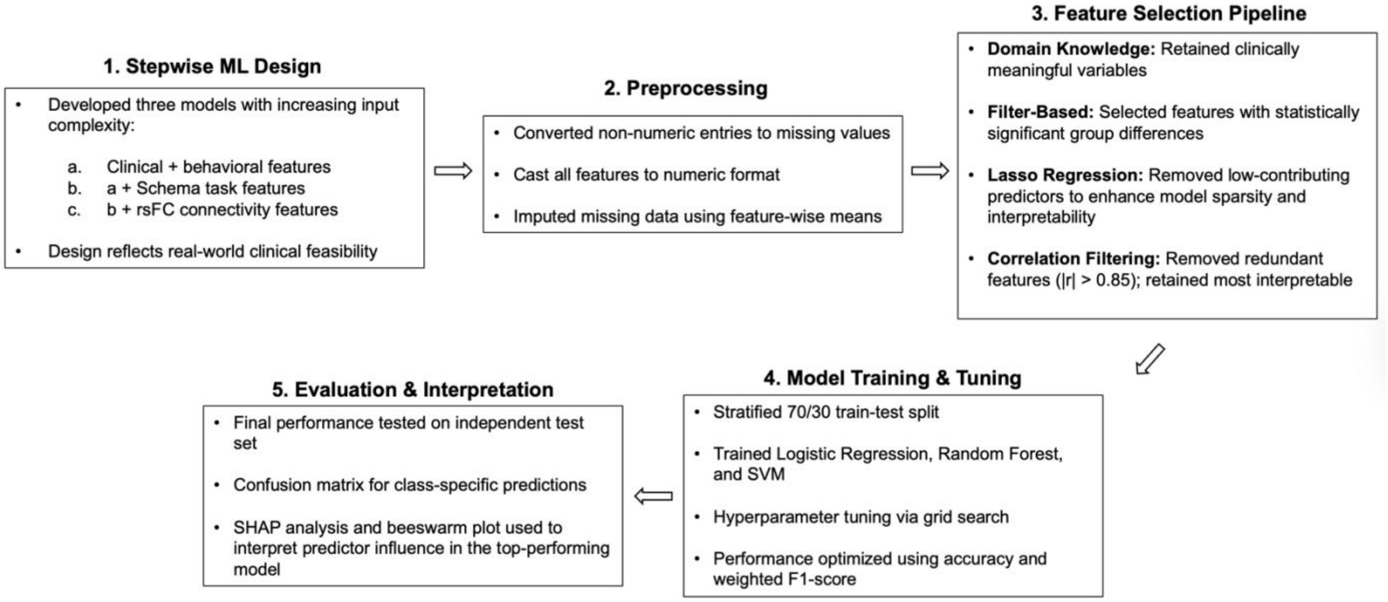

***Supplementary Figure 10.*** Overview of the supervised machine learning pipeline. The stepwise design progressed from clinical and behavioral features to schema task metrics and resting-state functional connectivity data. Each stage involved standardized preprocessing, feature selection using Lasso regression, correlation filtering, and model training with evaluation based on accuracy and weighted F1-score.

#### 6. Within-group analysis of matched vs mismatched vs uncued task events & group differences in uncued conditions

Friedman tests revealed significant effects at 47°C in all groups (High PDA: $\chi^2(2) = 55.088$, $p < 0.001$; Low PDA: $\chi^2(2) = 56.888$, $p < 0.001$; HC: $\chi^2(2) = 55.538$, $p < 0.001$), while at 45°C, significant differences were observed only in the High PDA ($\chi^2(2) = 10.091$, $p = 0.006$) and Low PDA groups ($\chi^2(2) = 18.586$, $p < 0.001$), with no effect in HC ($\chi^2(2) = 1.903$, $p = 0.386$). Wilcoxon Signed-Rank tests showed that in both high and low PDA groups, pain ratings in the mismatched condition were significantly higher than in the uncued condition at 47°C (High PDA: $Z = -6.257$, $p < 0.001$; Low PDA: $Z = -6.496$, $p < 0.001$) and 45°C (High PDA: $Z = -3.500$, $p < 0.001$; Low PDA: $Z = -3.959$, $p < 0.001$). In contrast, the HC group showed significant differences only at 47°C, where pain was higher in the mismatched condition than in the uncued condition ($Z = -6.147$, $p < 0.001$), and the matched condition also elicited higher ratings than the uncued condition ($Z = -2.495$, $p = 0.013$). No significant effects were found in HC at 45°C (all $p > 0.05$). Additionally, to examine whether pain ratings in the uncued conditions differed across groups, Kruskal–Wallis tests were conducted separately for 45°C and 47°C (see Supplementary Figure 11). At 45°C, no significant differences were observed between High PDA, Low PDA, and healthy control groups ($H(2) = 1.23$, $p = 0.541$). Similarly, at 47°C, the group effect did not reach significance ($H(2) = 3.93$, $p = 0.140$). Bonferroni-adjusted post-hoc comparisons revealed no significant pairwise differences between any groups at either temperature.

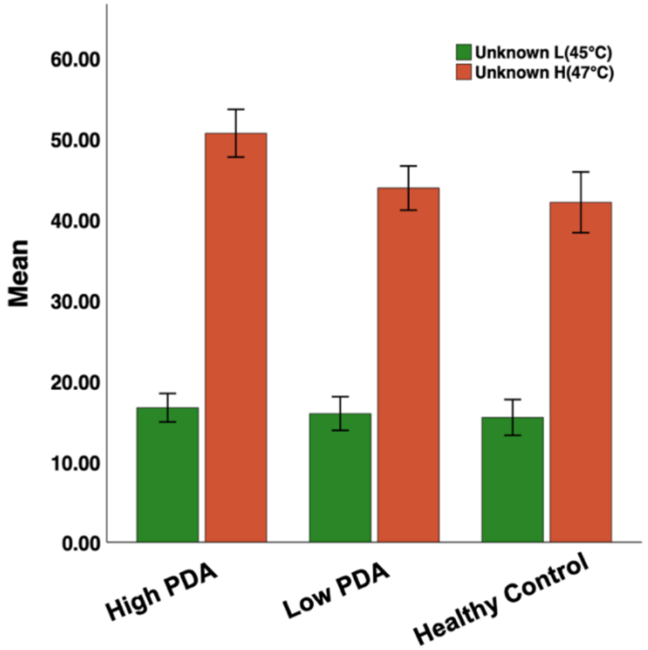

***Supplementary Figure 11.*** Mean pain ratings in uncued conditions (45°C and 47°C) across groups (High PDA, Low PDA, and Healthy Controls). Mean pain ratings for the uncued conditions, where no visual cue was provided prior to the heat stimulus, are shown for both uncued L (45°C) (green bars) and uncued H (47°C) (brown bars) conditions across the three groups: High PDA, Low PDA, and Healthy Controls. Error bars represent the standard error of the mean (SEM). While the 47°C condition consistently elicited higher pain ratings than the 45°C condition across all groups, independent-samples Kruskal-Wallis tests revealed no statistically significant differences in pain ratings between groups at either temperature: 45°C, $H(2) = 1.23$, $p = 0.541$; 47°C, $H(2) = 3.93$, $p = 0.140$.

#### 7. Group Differences in Top-Down Prediction Error Response

A Kruskal–Wallis test was conducted to compare the top-down prediction error response across groups. The analysis revealed no significant group differences ($H(2) = 1.76$, $p = 0.416$). The distribution of values is illustrated in Supplementary Figure 12.

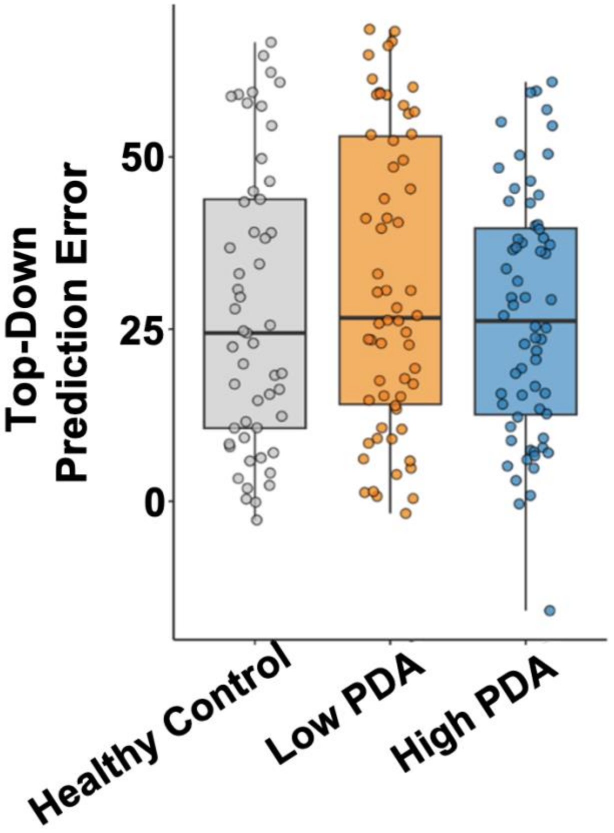

***Supplementary Figure 12.*** Top-Down Prediction Error response: Measures the maximum cognitive effect on pain perception by comparing pain ratings between the lowest and highest 20% of visual cue ranges, both linked to a 47°C stimulus, highlighting the maximal influence of expectations despite identical thermal inputs. No significant difference has been observed across the groups.

#### 8. Between Bottom-Up Prediction Error (and Max) and Affective Measures

Significant positive correlations were observed between bottom-up prediction error response and pain catastrophizing (PCS Total Score; $\rho = 0.148$, $p = 0.035$), as well as between bottom-up prediction error max and pain vigilance (PVAQ Total Score; $\rho = 0.198$, $p = 0.019$). These associations are visualized in Supplementary Figure 13.

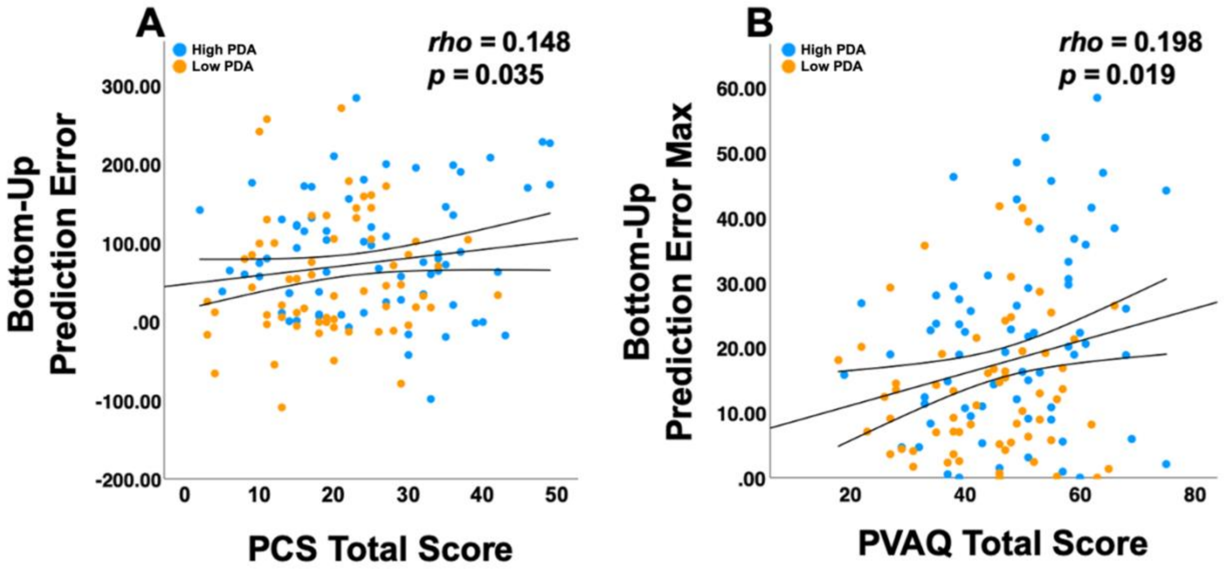

***Supplementary Figure 13.*** Significant correlations between prediction error bias metrics and behavioral measures in chronic pain populations. ***A,*** illustrates the positive correlation between prediction error bias and PCS Total Score, suggesting an association between threat bias and pain catastrophizing. ***B,*** shows the positive correlation between prediction error bias max and PVAQ Total Score, linking heightened threat sensitivity with increased pain vigilance. Data points are color-coded by group: blue represents the high PDA group, and orange represents the low PDA group. Trend lines indicate the direction and strength of each association. Abbreviations: PCS, Pain Catastrophizing Scale; PVAQ, Pain Vigilance and Awareness Questionnaire.

#### 9. Within-Group Connectivity Differences Between dl/lPAG and vlPAG (HC, high PDA, low PDA)

In the HC group, for the right hemisphere (Supplementary figure 14, Supplementary table 2), the right dl/lPAG exhibited stronger FC than the right vlPAG with regions primarily involved in salience and sensory processing, such as the right and left thalamus (Thal_R, Thal_L), the right intracalcarine cortex (IC_R), the right supramarginal gyrus, posterior division (SMGp_R), and the right and left inferior frontal gyrus, pars opercularis (IFGpo_R, IFGpo_L). Additional regions included the right inferior frontal gyrus, pars triangularis (IFGpt_L), the left rostral anterior cingulate cortex, mid-posterior division (ACCrm_L), and the left middle frontal gyrus (MFG_L). In contrast, the right vlPAG relative to the right dlPAG exhibited stronger FC with regions implicated in emotional and cognitive processing, including the right and left parahippocampal gyrus, posterior division (pHippp_R, pHippp_L), the right hippocampus (Hipp_R), the left posterior and middle insula (INSp_L, INSm_L), the left and right Heschl's gyrus (He_L, He_R), and the left planum polare (PlP_L).

For the left hemisphere (Supplementary figure 15, Supplementary table 3), the left dl/lPAG showed stronger FC than the left vlPAG with regions associated with salience and sensory integration, including the left and right thalamus (Thal_L, Thal_R), the left parietal operculum cortex (Pop_L), the left supramarginal gyrus, posterior and anterior divisions (SMGp_L, SMGa_L), the right and left inferior frontal gyrus, pars opercularis (IFGpo_R, IFGpo_L), the left inferior frontal gyrus, pars triangularis (IFGpt_L), and the left and right rostral anterior cingulate cortex, mid-posterior and posterior divisions (ACCrm_L, ACCrm_R, ACCrp_L, ACCrp_R). Additional regions included the right and left putamen (Put_R, Put_L). In contrast, the left vlPAG relative to the right dlPAG exhibited stronger FC with regions involved in emotional and memory processing, including the left and right parahippocampal gyrus, posterior division (pHippp_L, pHippp_R), and the right hippocampus (Hipp_R).

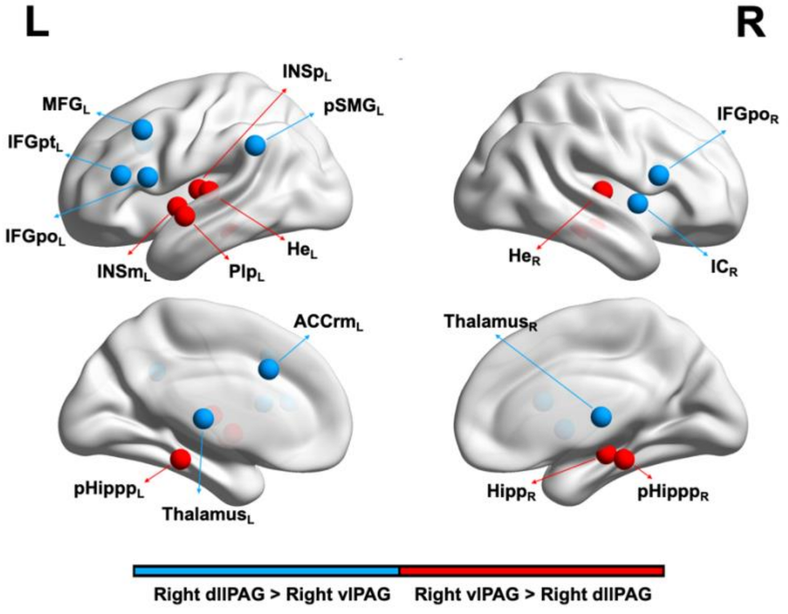

***Supplementary Figure 14.*** Resting-state functional connectivity (rsFC) for the right dl/lPAG and vlPAG within the healthy control (HC; n = 53) group. This figure illustrates the significant functional connectivity patterns (FDR-corrected, $p < 0.05$) for the right dl/lPAG compared to the right vlPAG. Blue arrows indicate regions where connectivity with the right dl/lPAG was stronger, while red arrows indicate regions where connectivity with the right vlPAG was stronger. ThalamusR/L, Right/Left thalamus; ICR, Right intracalcarine cortex; IFGpoR/L, Right/Left inferior frontal gyrus, pars opercularis; SMGpR, Right supramarginal gyrus, posterior division; IFGptL, Left inferior frontal gyrus, pars triangularis; MFGL, Left middle frontal gyrus; ACCrmL, Left rostral anterior cingulate cortex, mid-posterior division; pHipppL/R, Left/Right parahippocampal gyrus, posterior division; HippR, Right hippocampus; INSmL, Left middle insula; INSpL, Left posterior insula; HeL/R, Left/Right Heschl's gyrus; PIPL, Left planum polare; dl/lPAG, Dorsolateral/lateral periaqueductal gray; vlPAG, Ventrolateral periaqueductal gray.

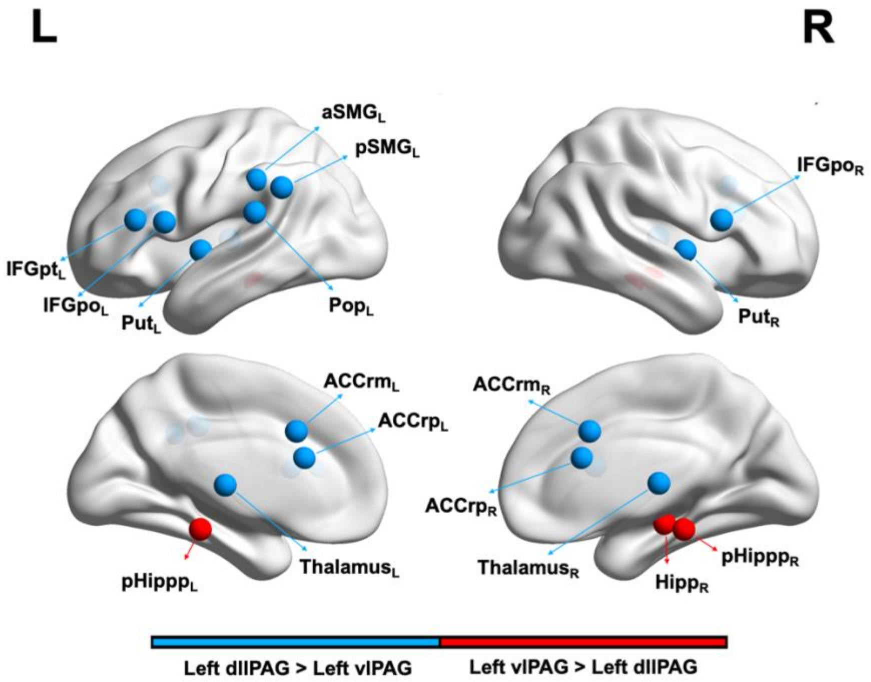

***Supplementary Figure 15.*** Resting-state functional connectivity (rsFC) for the left dl/lPAG and vlPAG within the healthy control (HC; n = 53) group. This figure illustrates the significant functional connectivity patterns (FDR-corrected, $p < 0.05$) for the left dl/lPAG compared to the left vlPAG. Blue arrows indicate regions where connectivity with the left dl/lPAG was stronger, while red arrows indicate regions where connectivity with the left vlPAG was stronger. ThalamusR/L, Right/Left thalamus; PopL, Left parietal operculum cortex; SMGpL, Left supramarginal gyrus, posterior division; SMGaL, Left supramarginal gyrus, anterior division; IFGpoR/L, Right/Left inferior frontal gyrus, pars opercularis; IFGptL, Left inferior frontal gyrus, pars triangularis; ACCrmR/L, Right/Left rostral anterior cingulate cortex, mid-posterior division; ACCrpR/L, Right/Left rostral anterior cingulate cortex, posterior division; PutR/L, Right/Left putamen; pHipppL/R, Left/Right parahippocampal gyrus, posterior division; HippR, Right hippocampus; dl/lPAG, Dorsolateral/lateral periaqueductal gray; vlPAG, Ventrolateral periaqueductal gray.

***Supplementary Table 2.*** Summary of significant functional connectivity findings for the right dl/lPAG and vlPAG within the healthy control group. Regions where the right dl/lPAG demonstrated stronger connectivity compared to the right vlPAG are listed in the first section, while regions with stronger connectivity to the right vlPAG are listed in the second section. For each region, the full name, MNI coordinates, Z-value, and p-value (with significance threshold $p < 0.05$) are provided.

| *Region of interest* | *MNI coordinates (x, y, z)* | *Z-value* | *P-value* |
|---|---|---|---|
| **Right dl/lPAG > Right vlPAG (Within HC)** | | | |

| | | | |
|---|---|---|---|
| R Thalamus (Thal_R) | 11, -18, 7 | 5.502 | <0.0001 |
| L Thalamus (Thal_L) | -10, -19, 6 | 5.4666 | <0.0001 |
| L Inferior frontal gyrus, pars opercularis (IFGpo_L) | -51 15 15 | 3.1383 | 0.0016 |
| L Supramarginal gyrus, posterior division (SMGp_L) | -55, -46, 33 | 3.0586 | 0.0022 |
| R Intracalcarine cortex (IC_R) | 37, 3, 0 | 2.7134 | 0.0066 |
| L Rostral anterior cingulate mid posterior (ACCrm_L) | -6, 18, 34 | 2.5806 | 0.0098 |
| R Inferior frontal gyrus, pars opercularis (IFGpo_R) | 52, 15, 16 | 2.5806 | 0.0098 |
| L Middle frontal gyrus (MFG_L) | -38, 18, 42 | 2.5363 | 0.0112 |
| L Inferior Frontal gyrus, pars triangularis (IFGpt_L) | -50, 30, 16 | 2.5186 | 0.0117 |
| **Right vlPAG > Right dl/lPAG (Within HC)** | | | |
| R Parahippocampal gyrus, posterior division (pHippp_R) | 23, -31, -17 | -5.5197 | <0.0001 |
| L Parahippocampal gyrus, posterior division (pHippp_L) | -22, -32, -17 | -4.4308 | <0.0001 |
| R Hippocampus (Hipp_R) | 26, -21, -14 | -3.218 | 0.0012 |
| L Posterior insula (INSp_L) | -38, -14, 8 | -3.0586 | 0.0022 |
| L Middle insula (INSm_L) | -40, -2, -2 | -2.917 | 0.0035 |
| L Heschl's gyrus (He_L) | -45, -20, 7 | -2.8019 | 0.005 |
| L Planum polare (PlP_L) | -47, -6, -7 | -2.6691, | 0.0076 |
| R Heschl's gyrus (He_R) | 46, -17, 7 | -2.5894 | 0.0096 |

***Supplementary Table 3.*** Summary of significant functional connectivity findings for the left dl/lPAG and vlPAG within the healthy control group. Regions where the left dl/lPAG demonstrated stronger connectivity compared to the left vlPAG are listed in the first section, while regions with stronger connectivity to the left vlPAG are listed in the second section. For each region, the full name, MNI coordinates, Z-value, and p-value (with significance threshold p < 0.05) are provided.

| ***Region of interest*** | ***MNI coordinates (x, y, z)*** | ***Z-value*** | ***P-value*** |
|---|---|---|---|
| **Left dl/lPAG > Left vlPAG (Within HC)** | | | |
| L Thalamus (Thal_L) | -10, -19, 6 | 5.7145 | <0.0001 |
| L Parietal operculum cortex (Pop_L) | -48, -32, 20 | 4.0059 | <0.0001 |
| L Supramarginal gyrus, posterior division (SMGp_L) | -55, -46, 33 | 3.9528 | <0.0001 |
| R Thalamus (Thal_R) | 11, -18, 7 | 3.9085 | <0.0001 |
| R Inferior frontal gyrus, pars opercularis (IFGpo_R) | 52, 15, 16 | 3.5367 | 0.0004 |
| L Inferior frontal gyrus, pars opercularis (IFGpo_L) | -51 15 15 | 3.218 | 0.0012 |
| L Supramarginal gyrus, anterior division (SMGa_L) | -57, -33, 37 | 3.0852 | 0.002 |
| L Rostral anterior cingulate posterior (ACCrp_L) | -4, 22, 20 | 3.0675 | 0.0021 |
| L Rostral anterior cingulate mid posterior (ACCrm_L) | -6, 18, 34 | 3.0498 | 0.0022 |

| | | | |
|---|---|---|---|
| R Putamen (Put_R) | 30, -4, 0 | 2.8373 | 0.0045 |
| R Rostral anterior cingulate mid posterior (ACCrm_R) | 6, 18, 34 | 2.8285 | 0.0046 |
| R Rostral anterior cingulate posterior (ACCrp_R) | 4, 22, 20 | 2.6071 | 0.0091 |
| L Inferior Frontal gyrus, pars triangularis (IFGpt_L) | -50, 30, 16 | 2.5629 | 0.0103 |
| L Putamen (Put_L) | -30, -4, 0 | 2.5009 | 0.0123 |
| **Left vlPAG > Left dl/lPAG (Within HC)** | | | |
| L Parahippocampal gyrus, posterior division (pHippp_L) | -22, -32, -17 | -5.8207 | <0.0001 |
| R Parahippocampal gyrus, posterior division (pHippp_R) | 23, -31, -17 | -4.4131 | <0.0001 |
| R Hippocampus (Hipp_R) | 26, -21, -14 | -2.855 | 0.0043 |

The connectivity profiles for the low and high PDA groups revealed both overlapping and distinct patterns relative to the HC group, highlighting variations in preferential rsFC between the dl/lPAG and vlPAG. In the low PDA group, several regions showing stronger connectivity with the dl/lPAG compared to the vlPAG in HC were reproduced, while others were absent or newly identified. Specifically, the connectivity with the rostral anterior cingulate cortex (ACCrm) and posterior rostral anterior cingulate cortex (ACCrp) observed in HC was absent in the low PDA group, whereas additional connectivity with regions such as the globus pallidus (GP), precuneus (PreC), precentral and postcentral gyri (PreC, PostC), superior colliculus (Sc), planum temporale (PlT), occipital pole (OccP), and frontal operculum (Fop) was detected. Conversely, regions showing stronger connectivity with the vlPAG in HC, such as the posterior insula (INSp) and Heschl's gyrus (He), were missing in the low PDA group, with additional connectivity emerging in areas including the middle temporal gyrus, anterior division (MTGa), posterior temporal fusiform cortex (TFCp), superior temporal gyrus, anterior division (STGa), anterior temporal fusiform cortex (TFCa), basolateral amygdala (BLA), and anterior ventral insula (vINSa).

In the high PDA group, similar trends were observed, with some findings overlapping with HC while others diverged. Connectivity with the ACCrm observed in HC was absent, while additional regions such as the cuneus (Cun), midcingulate cortex (ACCm), GP, Sc, and Fop displayed stronger connectivity with the dl/lPAG compared to the vlPAG. Similarly, regions showing stronger connectivity with the vlPAG in HC, including the INSp, middle insula (INSm), planum polare (PlP), and He, were absent in the high PDA group. Additional connectivity was identified in the MTGa, TFCp, vINSa, inferior temporal gyrus, anterior division (ITGa), BLA, and caudal anterior cingulate cortex (ACCc). All findings for each group are detailed in Supplementary Tables 4-7.

***Supplementary Table 4.*** Summary of significant functional connectivity findings for the right dl/lPAG and vlPAG within the high PDA group (n = 64). Regions where the right dl/lPAG demonstrated stronger connectivity compared to the left vlPAG are listed in the first section, while regions with stronger connectivity to the right vlPAG are listed in the second section. For each region, the full name, MNI coordinates, Z-value, and p-value (with significance threshold p < 0.05) are provided.

| *Region of interest* | *MNI coordinates (x, y, z)* | *Z-value* | *P-value* |
|---|---|---|---|
| **Right dl/lPAG > Right vlPAG (Within High PDA)** | | | |
| R Thalamus (Thal_R) | 11, -18, 7 | 6.0789 | <0.0001 |
| L Thalamus (Thal_L) | -10, -19, 6 | 5.3233 | <0.0001 |
| R Cuneal cortex (Cun_R) | 4, -82, 30 | 3.3103 | 0.0009 |
| L Cuneal cortex (Cun_L) | -4, -82, 30 | 3.21 | 0.0013 |
| R Supramarginal gyrus, posterior division (SMGp_R) | 55, -40, 34 | 2.9425 | 0.0032 |
| R Parietal operculum cortex (Pop_R) | 48, -32, 20 | 2.7887 | 0.0052 |
| R Supramarginal gyrus, anterior division (SMGa_R) | 57, -33, 37 | 2.5747 | 0.0103 |
| R Mid anterior cingulate (ACCm_R) | 6, -2, 42 | 2.4811 | 0.013 |
| L Globus Pallidus (GP_L) | -16, -2, -2 | 2.4476 | 0.0143 |
| R Globus Pallidus (GP_R) | 16, -2, -2 | 2.3874 | 0.0169 |
| **Right vlPAG > Right dl/lPAG (Within High PDA)** | | | |
| R Parahippocampal gyrus, posterior division (pHippp_R) | 23, -31, -17 | -5.9385 | <0.0001 |
| L Parahippocampal gyrus, posterior division (pHippp_L) | -22, -32, -17 | -5.2497 | <0.0001 |
| L Hippocampus (Hipp_L) | -26, -21, -14 | -4.3669 | <0.0001 |
| L Ventral anterior insula (vINSa_L) | -36, 10, -14 | -3.3906 | 0.0006 |
| L Parahippocampal gyrus, anterior division (pHippa_L) | -24, -6, -34 | -3.0963 | 0.0019 |
| R Hippocampus (Hipp_R) | 26, -21, -14 | -2.9425 | 0.0032 |
| R Parahippocampal gyrus, anterior division (pHippa_R) | 24, -6, -34 | -2.6148 | 0.0089 |
| R Temporal fusiform cortex, posterior division (TFCp_R) | 36, -16, -32 | -2.6014 | 0.0092 |
| L Middle temporal gyrus, anterior division (MTGa_L) | -58, -2, -22 | -2.4075 | 0.016 |

***Supplementary Table 5.*** Summary of significant functional connectivity findings for the left dl/lPAG and vlPAG within the high PDA group (n = 64). Regions where the left dl/lPAG demonstrated stronger connectivity compared to the left vlPAG are listed in the first section, while regions with stronger connectivity to the left vlPAG are listed in the second section. For each region, the full name, MNI coordinates, Z-value, and p-value (with significance threshold p < 0.05) are provided.

| *Region of interest* | *MNI coordinates (x, y, z)* | *Z-value* | *P-value* |
|---|---|---|---|
| **Left dl/lPAG > Left vlPAG (Within High PDA)** | | | |
| L Thalamus (Thal_L) | -10, -19, 6 | 5.3701 | <0.0001 |
| R Thalamus (Thal_R) | 11, -18, 7 | 5.0357 | <0.0001 |
| R Cuneal cortex (Cun_R) | 4, -82, 30 | 3.6714 | 0.0002 |

| | | | |
|---|---|---|---|
| R Mid anterior cingulate (ACCm_R) | 6, -2, 42 | 3.6179 | 0.0002 |
| R Parietal operculum cortex (Pop_R) | 48, -32, 20 | 3.5109 | 0.0004 |
| R Supramarginal gyrus, posterior division (SMGp_R) | 55, -40, 34 | 3.3438 | 0.0008 |
| R Supramarginal gyrus, anterior division (SMGa_R) | 57, -33, 37 | 3.2568 | 0.0011 |
| L Cuneal cortex (Cun_L) | -4, -82, 30 | 3.2368 | 0.0012 |
| L Supramarginal gyrus, anterior division (SMGa_L) | -57, -33, 37 | 3.2368 | 0.0012 |
| R Supracalcarine cortex (Sc_R) | 2, -84, 12 | 3.0763 | 0.002 |
| R Inferior Frontal gyrus, pars triangularis (IFGpt_R) | 50, 30, 16 | 3.0161 | 0.0025 |
| R Inferior frontal gyrus, pars opercularis (IFGpo_R) | 51, 15, 15 | 2.8623 | 0.0042 |
| R Globus Pallidus (GP_R) | 16, -2, -2 | 2.8422 | 0.0044 |
| R Putamen (Put_R) | 30, -4, 0 | 2.6817 | 0.0073 |
| L Globus Pallidus (GP_L) | -16, -2, -2 | 2.675 | 0.0074 |
| L Supramarginal gyrus, posterior division (SMGp_L) | -55, -46, 33 | 2.6349 | 0.0084 |
| L Rostral anterior cingulate posterior (ACCrp_L) | -4, 22, 20 | 2.6148 | 0.0089 |
| L Mid anterior cingulate (ACCm_L) | -6, -2, 42 | 2.5814 | 0.0098 |
| R Frontal operculum cortex (Fop_R) | 40, 20, 4 | 2.5346 | 0.0112 |
| R Rostral anterior cingulate posterior (ACCrp_R) | -4, 22, 20 | 2.4343 | 0.0149 |
| **Left vlPAG > Left dl/lPAG (Within High PDA)** | | | |
| L Parahippocampal gyrus, posterior division (pHippp_L) | -22, -32, -17 | -6.7276 | <0.0001 |
| L Hippocampus (Hipp_L) | -26, -21, -14 | -5.1895 | <0.0001 |
| R Parahippocampal gyrus, posterior division (pHippp_R) | 23, -31, -17 | -4.3937 | <0.0001 |
| R Hippocampus (Hipp_R) | 26, -21, -14 | -3.5912 | 0.0003 |
| L Parahippocampal gyrus, anterior division (pHippa_L) | -24, -6, -34 | -3.5845 | 0.0003 |
| R Parahippocampal gyrus, anterior division (pHippa_R) | 24, -6, -34 | -2.8957 | 0.0037 |
| L Middle temporal gyrus, posterior division (MTGp_L) | -62, -22, -18 | -2.7419 | 0.0061 |
| L Inferior temporal gyrus, anterior division (ITGa_L) | -50, -6, -40 | -2.5814 | 0.0098 |
| R Inferior temporal gyrus, anterior division (ITGa_R) | 50, -6, -40 | -2.5011 | 0.0123 |
| L Caudal anterior cingulate (ACCc_L) | -4, 40, -2 | -2.3473 | 0.0189 |
| R Basolateral Amygdala (BLA_R) | 24, -2, -20 | -2.3139 | 0.0206 |

***Supplementary Table 6.*** Summary of significant functional connectivity findings for the right dl/lPAG and vlPAG within the low PDA group (n = 62). Regions where the right dl/lPAG demonstrated stronger connectivity compared to the left vlPAG are listed in the first section, while regions with stronger connectivity to the right vlPAG are listed in the second section. For each region, the full name, MNI coordinates, Z-value, and p-value (with significance threshold p < 0.05) are provided.

| *Region of interest* | *MNI coordinates (x, y, z)* | *Z-value* | *P-value* |
|---|---|---|---|
| **Right dl/lPAG > Right vlPAG (Within Low PDA)** | | | |

| | | | |
|---|---|---|---|
| R Thalamus (Thal_R) | 11, -18, 7 | 6.2293 | <0.0001 |
| L Thalamus (Thal_L) | -10, -19, 6 | 5.7245 | <0.0001 |
| L Supramarginal gyrus, posterior division (SMGp_L) | -55, -46, 33 | 3.7895 | 0.0001 |
| R Putamen (Put_R) | 30, -4, 0 | 3.6703 | 0.0002 |
| L Putamen (Put_L) | -30, -4, 0 | 3.5301 | 0.0004 |
| R Globus Pallidus (GP_R) | 16, -2, -2 | 3.3688 | 0.0007 |
| R Parietal operculum cortex (Pop_R) | 48, -32, 20 | 3.1725 | 0.0015 |
| R Inferior frontal gyrus, pars opercularis (IFGpo_R) | 51 15 15 | 3.1304 | 0.0017 |
| L Globus Pallidus (GP_L) | -16, -2, -2 | 3.1024 | 0.0019 |
| L Supramarginal gyrus, anterior division (SMGa_L) | -57, -33, 37 | 2.9552 | 0.0031 |
| R Supramarginal gyrus, posterior division (SMGp_R) | 55, -40, 34 | 2.8851 | 0.0039 |
| L Inferior frontal gyrus, pars opercularis (IFGpo_L) | -51 15 15 | 2.829 | 0.0046 |
| R Supramarginal gyrus, anterior division (SMGa_R) | 57, -33, 37 | 2.822 | 0.0047 |
| L Precentral gyrus (PreC_L) | -44, -8, 52 | 2.7939 | 0.0052 |
| R Precentral gyrus (PreC_R) | 44, -8, 52 | 2.7238 | 0.0064 |
| R Intracalcarine cortex (IC_R) | 37, 3, 0 | 2.7028 | 0.0068 |
| R Planum temporale (PlT_R) | 60, -22, 8 | 2.6747 | 0.0074 |
| R Supracalcarine cortex (Sc_R) | 2, -84, 12 | 2.5485 | 0.0108 |
| R Occipital pole (OccP_R) | 8, -100, 6 | 2.4153 | 0.0157 |
| L Occipital pole (OccP_L) | -8, -100, 6 | 2.3662 | 0.0179 |
| R Inferior Frontal gyrus, pars triangularis (IFGpt_R) | 50, 30, 16 | 2.3592 | 0.0183 |
| L Inferior Frontal gyrus, pars triangularis (IFGpt_L) | -50, 30, 16 | 2.3101 | 0.0208 |
| R Postcentral gyrus (PostC_R) | 50, -20, 46 | 2.1629 | 0.0305 |
| **Right vlPAG > Right dl/lPAG (Within Low PDA)** | | | |
| R Parahippocampal gyrus, posterior division (pHippp_R) | 23, -31, -17 | -5.4651 | <0.0001 |
| L Parahippocampal gyrus, posterior division (pHippp_L) | -22, -32, -17 | -5.3039 | <0.0001 |
| L Parahippocampal gyrus, anterior division (pHippa_L) | -24, -6, -34 | -4.1891 | <0.0001 |
| R Parahippocampal gyrus, anterior division (pHippa_R) | 24, -6, -34 | -3.8105 | 0.0001 |
| R Middle temporal gyrus, anterior division (MTGa_R) | 58, -2, -22 | -3.6913 | 0.0002 |
| L Hippocampus (Hipp_L) | -26, -21, -14 | -3.502 | 0.0004 |
| L Planum polare (PlP_L) | -47, -6, -7 | -3.3688 | 0.0007 |
| R Temporal fusiform cortex, posterior division (TFCp_R) | 36, -16, -32 | -3.2426 | 0.0011 |
| R Superior temporal gyrus, anterior division (STGa_R) | 58, -4, -6 | -2.9622 | 0.003 |
| L Middle temporal gyrus, posterior division (MTGp_L) | -62, -22, -18 | -2.9341 | 0.0033 |
| L Superior temporal gyrus, anterior division (STGa_L) | -58, -4, -6 | -2.6397 | 0.0082 |
| R Temporal fusiform cortex, anterior division (TFCa_R) | 32, -6, -42 | -2.5415 | 0.011 |
| L Temporal fusiform cortex, anterior division (TFCa_L) | -32, -6, -42 | -2.4784 | 0.0131 |
| L Middle temporal gyrus, anterior division (MTGa_L) | -58, -2, -22 | -2.4293 | 0.0151 |
| L Basolateral Amygdala (BLA_L) | -24, -2, -20 | -2.3172 | 0.0204 |
| L Middle insula (INSm_L) | -40, -2, -2 | -2.1769 | 0.0294 |

***Supplementary Table 7.*** Summary of significant functional connectivity findings for the left dl/lPAG and vlPAG within the low PDA group (n = 62). Regions where the left dl/lPAG demonstrated stronger connectivity compared to the left vlPAG are listed in the first section, while regions with stronger connectivity to the left vlPAG are listed in the second section. For each region, the full name, MNI coordinates, Z-value, and p-value (with significance threshold p < 0.05) are provided.

| ***Region of interest*** | ***MNI coordinates (x, y, z)*** | ***Z-value*** | ***P-value*** |
|---|---|---|---|
| **Left dl/lPAG > Left vlPAG (Within Low PDA)** | | | |
| L Thalamus (Thal_L) | -10, -19, 6 | 6.7972 | <0.0001 |
| R Thalamus (Thal_R) | 11, -18, 7 | 6.3135 | <0.0001 |
| L Putamen (Put_L) | -30, -4, 0 | 5.1286 | <0.0001 |
| R Putamen (Put_R) | 30, -4, 0 | 4.6589 | <0.0001 |
| L Inferior frontal gyrus, pars opercularis (IFGpo_L) | -51, 15, 15 | 4.5607 | <0.0001 |
| L Inferior Frontal gyrus, pars triangularis (IFGpt_L) | -50, 30, 16 | 4.3363 | <0.0001 |
| R Globus Pallidus (GP_R) | 16, -2, -2 | 4.2172 | <0.0001 |
| R Parietal operculum cortex (Pop_R) | 48, -32, 20 | 3.7685 | 0.0001 |
| R Inferior frontal gyrus, pars opercularis (IFGpo_R) | 51, 15, 15 | 3.7614 | 0.0001 |
| L Supramarginal gyrus, posterior division (SMGp_L) | -55, -46, 33 | 3.6493 | 0.0002 |
| R Inferior Frontal gyrus, pars triangularis (IFGpt_R) | 50, 30, 16 | 3.5161 | 0.0004 |
| L Globus Pallidus | -16, -2, -2 | 3.2707 | 0.001 |
| L Supramarginal gyrus, anterior division (SMGa_L) | -57, -33, 37 | 3.2006 | 0.0013 |
| R Precentral gyrus (PreC_R) | 44, -8, 52 | 3.1935 | 0.0014 |
| R Frontal operculum cortex (Fop_R) | 40, 20, 4 | 3.1865 | 0.0014 |
| R Intracalcarine cortex (IC_R) | 37, 3, 0 | 2.9411 | 0.0032 |
| L Precentral gyrus (PreC_L) | -44, -8, 52 | 2.9131 | 0.0035 |
| L Frontal operculum cortex (Fop_L) | -40, 20, 4 | 2.7939 | 0.0052 |
| R Supramarginal gyrus, anterior division (SMGa_R) | 57, -33, 37 | 2.7308 | 0.0063 |
| R Dorsal anterior insula (dINSa_R) | 32, 20, 0 | 2.6537 | 0.0079 |
| R Supramarginal gyrus, posterior division (SMGp_R) | 55, -40, 34 | 2.6397 | 0.0082 |
| L Middle frontal gyrus (MFG_L) | -40, 20, 44 | 2.5275 | 0.0114 |
| R Planum temporale (PlT_R) | 60, -22, 8 | 2.4854 | 0.0129 |
| R Supplementary motor area (SMA_R) | 4, -2, 58 | 2.4434 | 0.0145 |
| L Parietal operculum cortex (Pop_L) | -48, -32, 20 | 2.4013 | 0.0163 |
| R Supracalcarine cortex (Sc_R) | 2, -84, 12 | 2.3522 | 0.0186 |
| R Central opercular cortex (Cop_R) | 48, -4, 8 | 2.3312 | 0.0197 |
| L Occipital pole (OccP_L) | -8, -100, 6 | 2.2821 | 0.0224 |
| **Left vlPAG > Left dl/lPAG (Within Low PDA)** | | | |
| L Parahippocampal gyrus, posterior division (pHippp_L) | -22, -32, -17 | -6.2854 | <0.0001 |
| R Parahippocampal gyrus, posterior division (pHippp_R) | 23, -31, -17 | -4.4415 | <0.0001 |
| L Parahippocampal gyrus, anterior division (pHippa_L) | -24, -6, -34 | -4.112 | <0.0001 |
| R Parahippocampal gyrus, anterior division (pHippa_R) | 24, -6, -34 | -3.7755 | 0.0001 |
| L Hippocampus (Hipp_L) | -26, -21, -14 | -3.3548 | 0.0007 |
| L Ventral anterior insula (vINSa_L) | -36, 10, -14 | -3.0463 | 0.0023 |

| | | | |
|---|---|---|---|
| L Planum polare (PlP_L) | -47, -6, -7 | -2.9271 | 0.0034 |
| L Basolateral Amygdala (BLA_L) | -24, -2, -20 | -2.8851 | 0.0039 |
| R Middle temporal gyrus, anterior division (MTGa_R) | 58, -2, -22 | -2.7659 | 0.0056 |
| R Temporal fusiform cortex, posterior division (TFCp_R) | 36, -16, -32 | -2.6817 | 0.0073 |
| L Temporal fusiform cortex, posterior division (TFCp_L) | -36, -16, -32 | -2.5135 | 0.0119 |

#### 10. dl/lPAG Connectivity: Group Differences and Correlations

Resting-state analyses showed that the high PDA group had significantly more negative dl/lPAG connectivity with multiple prefrontal and limbic regions compared to healthy controls. We then compared high and low PDA groups, limiting the analysis to regions identified in the high PDA vs. HC contrast. Within this subset, connectivity between the dl/lPAG and bilateral dmPFCa remained significantly more negative in the high PDA group (see Supplementary Table 8). Supplementary Figure 16 displays additional correlations not included in the main text, showing that reduced dl/lPAG–dmPFCa connectivity was associated with heightened bottom-up prediction error responses.

***Supplementary Table 8.*** This table summarizes regions exhibiting significant differences in resting-state functional connectivity (rsFC) with the left and right dl/lPAG. The first section presents results from the comparison between high PDA and Healthy Controls, showing all significant clusters for both hemispheres. The second section presents differences between high and low PDA groups, restricted to regions previously identified in the comparison with Healthy Controls. For each region, the anatomical label, MNI coordinates (x, y, z), U-statistic, and p-value are reported. All results are significant after FDR correction ($p < 0.05$).

| Seed Region | Group Comparison | Region Name | MNI Coordinates (x, y, z) | U | p-value |
|---|---|---|---|---|---|
| ***Left dl/lPAG*** | ***High PDA vs. HC*** | Left dMPFCa | -4, 50, 28 | 3085.00 | 0.0002 |
| ***Right dl/lPAG*** | ***High PDA vs. HC*** | Left dMPFCa | -4, 50, 28 | 2873.00 | 0.0001 |
| | | Right dMPFCa | 4, 50, 28 | 2930.00 | 0.0001 |
| | | Left ACCr | -4, 38, 18 | 3101.00 | 0.0002 |
| | | Left NAcC | -12, 10, -8 | 3113.00 | 0.0003 |
| | | Left dMPFCp | -4, 26, 48 | 3136.00 | 0.0005 |
| | | Left MPFC | -6, 60, 8 | 3172.00 | 0.0009 |
| | | Right ACCr | 4, 38, 18 | 3198.00 | 0.0016 |
| | | Left FP | -30, 54, 20 | 3220.00 | 0.0023 |
| | | Right NAcS | 9, 10, -6 | 3240.00 | 0.0033 |
| | | Right dMPFCp | 4, 26, 48 | 3250.00 | 0.004 |
| | | Right MPFC | 6, 60, 8 | 3263.00 | 0.0044 |
| ***Right dl/lPAG*** | ***High PDA vs. Low PDA*** | Left dMPFCa | -4, 50, 28 | 3383.00 | 0.0009 |
| | | Right dMPFCa | 4, 50, 28 | 3482.00 | 0.0045 |

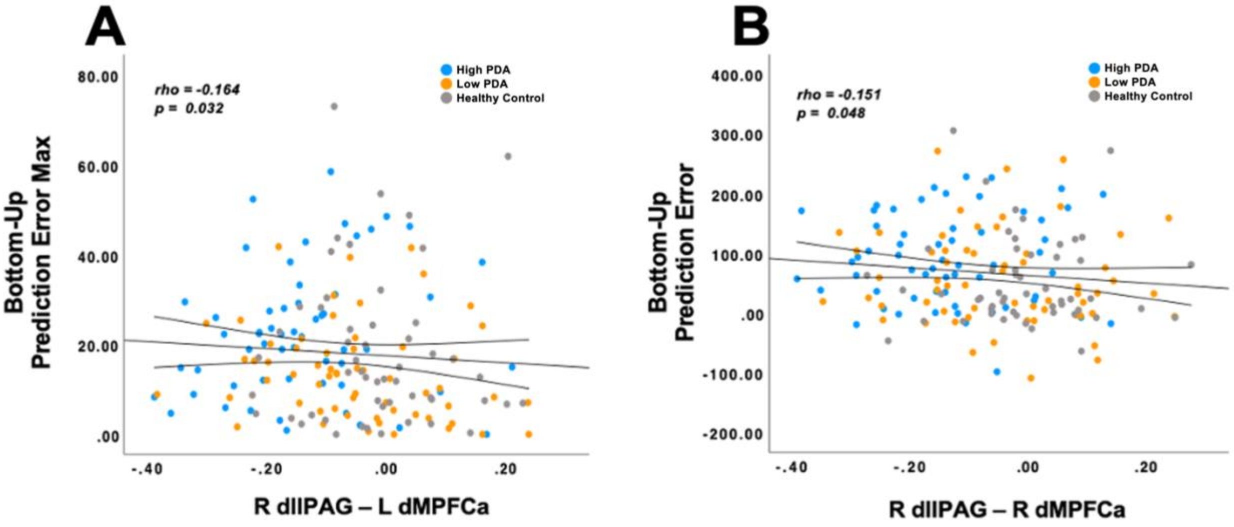

***Supplementary Figure 16.*** Correlations between schema-derived prediction error metrics and targeted seed-based PAG connectivity. Scatterplots show significant negative associations between dl/lPAG–dmPFCa resting-state connectivity and pain modulation metrics derived from the schema task. (A) Connectivity between the right dl/lPAG and left dmPFCa was negatively correlated with bottom-up prediction error max ($\rho = -0.164$, $p = 0.032$). (B) Connectivity between the right dl/lPAG and right dmPFCa was negatively correlated with bottom-up prediction error ($\rho = -0.151$, $p = 0.048$). Data points are color-coded by group: blue = High PDA, orange = Low PDA, gray = HC. These results suggest that reduced functional coupling between the dl/lPAG and dmPFCa tracks with amplified expectation-driven prediction error responses, particularly in individuals with high PDA. dl/lPAG = dorsolateral/lateral periaqueductal gray; dmPFCa = dorsomedial prefrontal cortex, anterior division; rsFC = resting-state functional connectivity.

#### 11. Resting-State Connectivity Analysis with Physiological Noise Correction (Supplementary)

To assess whether physiological noise contributed to individual differences in resting-state functional connectivity (rsFC), we conducted a supplementary analysis incorporating peripheral physiological recordings. Respiration was measured using a belt placed around the torso, and cardiac signals were recorded using a photoplethysmography (PPG) sensor attached to the finger. These recordings were available for a subset of participants in the high and low Pain–Disability–Affect (PDA) groups; healthy controls were excluded due to missing data.

Physiological regressors were generated using RETROICOR modeling in the PhysIO Toolbox (Kasper et al., 2017) and included in an updated nuisance regression model, alongside six motion parameters, cerebrospinal fluid (CSF), white matter (WM), and global signal. The remaining preprocessing steps, including motion correction, distortion correction, spatial smoothing, temporal filtering, and registration, were identical to those used in the primary analysis.

Following nuisance regression, BOLD time series were re-extracted from all 134 atlas-defined regions using the Optimized Harvard-Oxford parcellation, as well as from the four PAG subregions (right/left dl/lPAG and vlPAG) defined in the main analysis. Pearson correlation coefficients were computed between each PAG seed and all 134 regions, resulting in 4 × 134 rsFC matrices per participant. Between-group comparisons were conducted using Mann–Whitney U tests, and false discovery rate (FDR) correction ($q < 0.05$) was applied across all tests.

In a follow-up analysis that compared high and low PDA groups within regions previously found to differ between high PDA participants and healthy controls, we observed both replicated and new findings using physiologically denoised data. Connectivity between the right dl/lPAG and the right and left dorsomedial prefrontal cortex (dmPFCa) remained significantly more reduced in the high PDA group, consistent with the original results (Supplementary Figure 17).

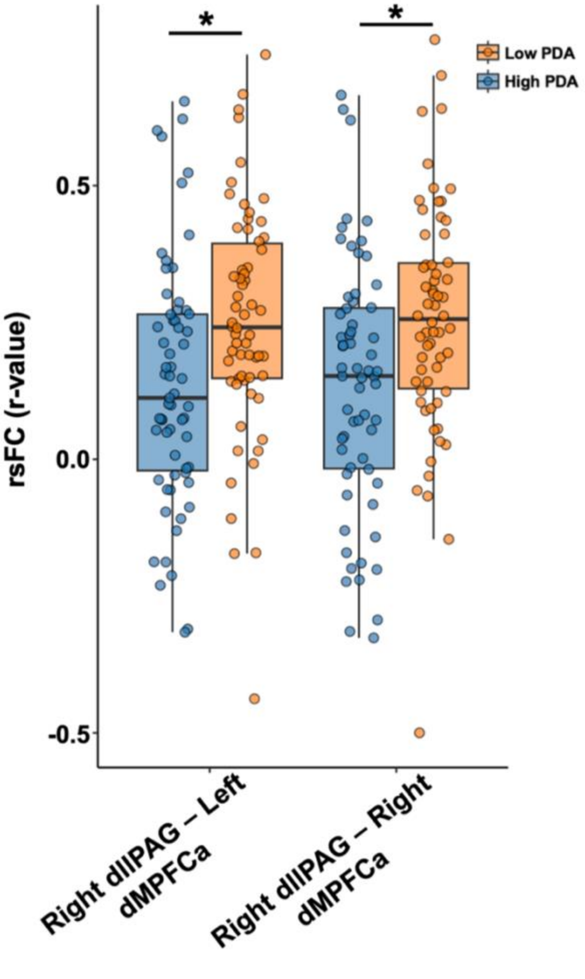

***Supplementary Figure 17.*** Resting-state functional connectivity between the right dl/lPAG and bilateral dmPFCa after physiological noise correction. The high PDA group (blue) showed significantly reduced connectivity relative to the low PDA group (orange), consistent with the original analysis (Mann–Whitney U tests, $p < 0.05$, FDR-corrected). Box plots display median, interquartile range, and individual data points. dl/lPAG, dorsolateral/lateral periaqueductal gray; dmPFCa, anterior dorsomedial prefrontal cortex; PDA, Pain–Disability–Affect.

#### 12. Logistic Regression and ROC Analysis for High vs. Low PDA Group Classification

To evaluate the utility of schema-based and rsFC features in distinguishing high from low PDA subgroups, receiver operating characteristic (ROC) analyses and logistic regression were conducted (Supplementary Figure 18). While individual metrics showed modest classification performance, a binary logistic regression using backward stepwise selection retained Bottom-Up Prediction Error Max and right dl/lPAG–left dmPFCa connectivity as significant predictors (Supplementary Table 9). The final model explained 24.7% of the variance (Nagelkerke $R^2$ = 0.247) and achieved 69.2% overall classification accuracy, with sensitivity of 68.4% and specificity of 70.0%, outperforming individual predictors.

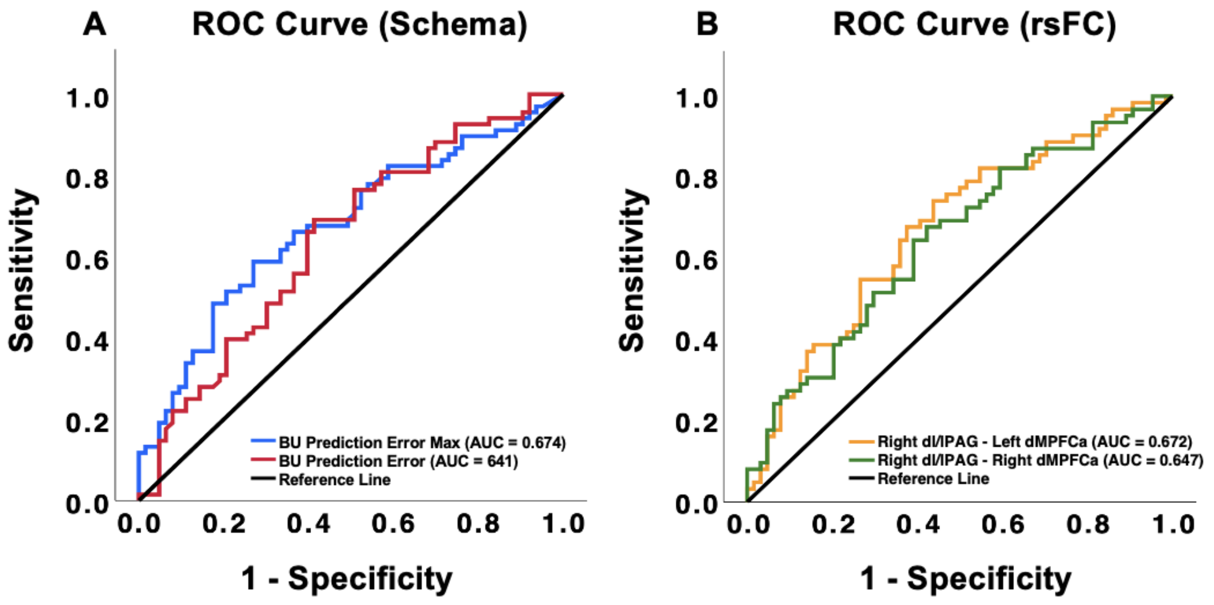

***Supplementary Figure 18.*** ROC curves depicting the predictive performance of schema-based and rs-FC metrics for chronic pain severity. ***A,*** Schema metrics show moderate discrimination, with Bottom-Up Prediction Error Max (AUC = 0.674) and Bottom-Up Prediction Error (AUC = 0.641). ***B,*** rs-FC metrics demonstrate stronger predictive utility, particularly for the low PDA group. Connectivity between the right dl/lPAG and left dMPFCa (AUC = 0.672) and right dMPFCa (AUC = 0.647) differentiates high and low PDA groups. The diagonal reference line represents random classification performance. dl/lPAG, dorsolateral/lateral periaqueductal gray; dMPFCa, dorsomedial prefrontal cortex anterior division; rs-FC, resting-state functional connectivity; IFGpo, inferior frontal gyrus pars opercularis; IFGpt, inferior frontal gyrus pars triangularis.

***Supplementary Table 9.*** Predictors retained in the final step of binary logistic regression. The table presents schema-related and resting-state functional connectivity (rs-FC) metrics identified as significant predictors in the final step of the backward stepwise logistic regression model. Coefficients (B), standard errors (SE), Wald statistics, p-values, odds ratios (Exp(B)), and 95% confidence intervals (CIs) for Exp(B) are shown. dl/lPAG, dorsolateral/lateral periaqueductal gray; dMPFCa, dorsomedial prefrontal cortex anterior division.

| Variable | B | S.E. | Wald | df | Sig. | Exp(B) | 95% CI for Exp(B) |
|---|---|---|---|---|---|---|---|
| **BU Prediction Error Max** | 0.054 | 0.017 | 9.946 | 1 | 0.002 | 1.055 | [1.021, 1.091] |
| **Right dl/lPAG – Left dMPFCa** | -5.079 | 1.637 | 9.625 | 1 | 0.002 | 0.006 | [0.000, 0.151] |
| **Constant** | -1.521 | 0.419 | 13.193 | 1 | <0.001 | 0.218 | |
